# Cellular and functional dissection of the octopaminergic system in the *Drosophila* brain

**DOI:** 10.64898/2026.02.06.704492

**Authors:** Lesly M. Palacios Castillo, Valentina Fajner, Ezgi Yalbir, Shafana Shahul, Bruce Ruff, Victor Junmyung Lee, Aundrea Koger, Juliet Heller, Kenichi Ishii, Veronica Morad, Max Trask, Carter Warren, Giovanni Frighetto, Mark Frye, Kenta Asahina

## Abstract

Octopamine (OA) is a major biogenic amine in the invertebrate nervous system and is often considered a functional analog of vertebrate noradrenaline. Along with its immediate precursor tyramine (TA), OA influences diverse physiological and behavioral processes, including sensory processing and social behavior. However, understanding the neural basis of its multifunctionality has been constrained by the limited genetic access to defined OA/TA neuron types. Here, we present a curated set of transgenic driver strains that provide selective access to nearly all long-range OA/TA cell types in the brain of common fruit flies, *Drosophila melanogaster*. Using these tools, we map cell-type-specific innervation patterns, compare male and female neuroanatomy, and cross-reference identified neuron types with electron microscopy connectome datasets. As a proof of principle, we show that distinct optic lobe-projecting OA/TA neuron types differentially modulate visually guided behaviors, and we identify a novel OA/TA cell type that suppresses aggression in both sexes. This resource establishes a practical and conceptual foundation for cell-type-resolved analysis of OA/TA circuit function and enables direct integration of genetics, anatomy, and connectomics for studies of neuromodulatory circuit organization.

## Introduction

Octopamine (OA) is an invertebrate-specific biogenic amine neuromodulator (Roeder, 2005), structurally similar to noradrenaline in vertebrates. High sequence similarity between vertebrate adrenergic receptors and invertebrate OA receptors (Evans and Maqueira, 2005) suggest that the physiological effects of OA and noradrenaline are similar at the cellular level. This similarity extends to neuroanatomical organization and behavior. Although adrenergic and octopaminergic cells are relatively few and localized within the central brain, some innervate extensively throughout the brain and even project to motor centers of the ventral nerve cord (Busch et al., 2009; Chandler et al., 2019). The long-range projection patterns of adrenergic neurons are well suited to modulate complex, multi-compartmental cognitive functions such as attention and arousal (Chandler et al., 2019; Schwarz and Luo, 2015). Studies in both mammals and insects indicate that noradrenaline and OA are involved in a variety of physiological and behavioral processes, including wakefulness, sensory modulation, learning, and memory. While such functional diversity is consistent with the high-level role classically attributed to the noradrenergic system, neuronal and cell circuit-based mechanisms through which a small population of noradrenergic or octopaminergic neurons can exert specialized, disparate, and occasionally opposing effects remain largely unexplored.

In insects, OA is synthesized from its immediate precursor tyramine (TA), a biogenic amine that also functions as a neuromodulator (Roeder, 2005). Tyrosine decarboxylase 2 (*Tdc2*), which converts tyrosine to tyramine, is a molecular marker of neurons that release either TA, OA, or both. Hereafter, neurons that express *Tdc2* are collectively referred to as OA/TA neurons. There are ∼70 OA/TA neurons with long-range projections in the *Drosophila* brain (Busch et al., 2009), in addition to ∼100 OA/TA neurons with relatively short-range restricted projection patterns (Wolff et al., 2025). Clonal analysis of *Tdc2*-expressing neurons revealed that the long-range OA/TA neurons consist of at least 28 anatomically distinct cell types from 4 major OA-immunoreactive clusters (Busch et al., 2009; Busch and Tanimoto, 2010). These neurons send distinct, far-reaching projections to multiple neuropils, collectively tiling the entire brain in a partially overlapping manner. This neuroanatomical heterogeneity within OA/TA neurons suggests that the diverse physiological and behavioral effects of OA/TA emerge from functionally distinct OA/TA neuronal subtypes, each with specialized functions. However, the lack of methods to systematically manipulate individual subtypes has left only fragmentary evidence in support of this hypothesis. Consequently, the field has been often constrained to treat the full ensemble of *Tdc2*-expressing neurons as a single functional unit.

The versatility of genetic tools and the scalability of behavioral assays in *Drosophila* make it a particularly powerful model to determine the molecular and cellular mechanisms of octopaminergic neuromodulation. In particular, the split-GAL4 technique (Luan et al., 2006; Meissner et al., 2025) enables genetic access to very small subsets of neurons, sometimes down to a single homogeneous bilateral pair, by subdividing otherwise heterogeneous driver lines. Large-scale screening efforts have identified comprehensive sets of split-GAL4 lines that cover many of the cell types interconnecting specific neuropils (Meissner et al., 2025), including the mushroom bodies, antennal lobe, lateral horn (Aso et al., 2014; Dolan et al., 2019; Shuai et al., 2025), optic lobes (Nern et al., 2025), central complex (Wolff et al., 2025; Wolff and Rubin, 2018), subesophageal zone (Sterne et al., 2021), as well as descending (Namiki et al., 2018; Zung et al., 2025) pathways. The relatively small number and well-defined anatomy of OA/TA neurons, combined with versatile genetic tools, are well suited to generate highly selective genetic reagents that allow cell type-specific functional characterization of these neurons.

In this study, we engineered and characterized a novel set of split-GAL4 lines that, together with some previously characterized split-GAL4 lines, label nearly all the known long-range OA/TA subtypes. These split-GAL4 lines show comparable expression patterns between the two sexes, consistent with previous reports that OA/TA neurons do not express sex-determining genes aside from a few exceptions (Andrews et al., 2014; Certel et al., 2010). Most cell types are labeled by more than one split-GAL4 line, enhancing experimental flexibility and reproducibility. In-depth examination of these split-GAL4 lines revealed several previously uncharacterized OA/TA cell types. Lastly, we demonstrate the power of this new tool by identifying specific OA/TA subtypes that suppresses aggression in both males and females, and by characterizing subtype-specific effects of octopaminergic modulation of visually guided flight behaviors, consistent with connectome-based predictions. Our work lays a foundation for comprehensively determining octopaminergic neuromodulation at single-cell and single-circuit resolution in the *Drosophila* brain, providing general mechanistic insights into the cellular principles underlying noradrenergic neuromodulation across nervous systems.

## Results

### Overview of the split-GAL4 lines for OA/TA neurons

To generate genetic reagents that enable the comprehensive physiological and behavioral characterization of OA/TA neurons, we examined 529 driver line combinations that were expected to label specific subtypes of OA/TA neurons (see Materials and Methods for the approaches used to choose hemi drivers). After initial screening, selected split-GAL4 lines with a reasonably high specificity were further characterized with anti-Tdc2 immunoreactivity, testing for expression consistency in brains of both sexes, and where necessary, clonal analysis of single cell type characterization using multi-color FLP-out (MCFO) technique (Nern et al., 2015). Here we report expression patterns of 39 lines, each of which labels three or fewer OA/TA neuronal classes in a complementary manner (Supplementary Table S1-3).

In-depth annotation of the *Drosophila* brain EM volumes (FAFB (Dorkenwald et al., 2024; Schlegel et al., 2024), MCNS (Berg et al., 2025), BANC (Bates et al., 2025)) enabled direct comparison of previously characterized OA/TA neuronal subtypes (Busch et al., 2009), with putative OA/TA neurons in these datasets, and with confocal images of our split-GAL4 lines. The number of OA/TA neurons identified on the Flywire/hemibrain platform (Dorkenwald et al., 2022; Scheffer et al., 2020; Schlegel et al., 2024) closely match with the number of *Tdc2-GAL4* neurons across the four OA/TA neuronal clusters (Table 1), based on cell body positions within the brain neuropils: anterior superior medial protocerebrum (ASM), antenna lobe (AL), ventral midline (VM), ventrolateral protocerebrum (VL) (Busch et al., 2009). In all four clusters, we detected consistent co-labeling of Tdc2 and GFP driven by the GAL4-expressing allele of the *vesicular monoamine transporter* (*Vmat*) gene (Supplementary Fig. S1A), which is necessary for packaging monoamines to neuronal vesicles. In addition, OA/TA neurons are co-labeled by *VGlut^LexA^*(Supplementary Fig. S1B), but not by *ChAT^LexA^* (Supplementary Fig. S1C) or by *GAD1^LexA^* (Supplementary Fig. S1D), consistent with previous reports that OA/TA neurons release glutamate as a co-transmitter (Croset et al., 2018; Sherer et al., 2020).

**Table 1:**
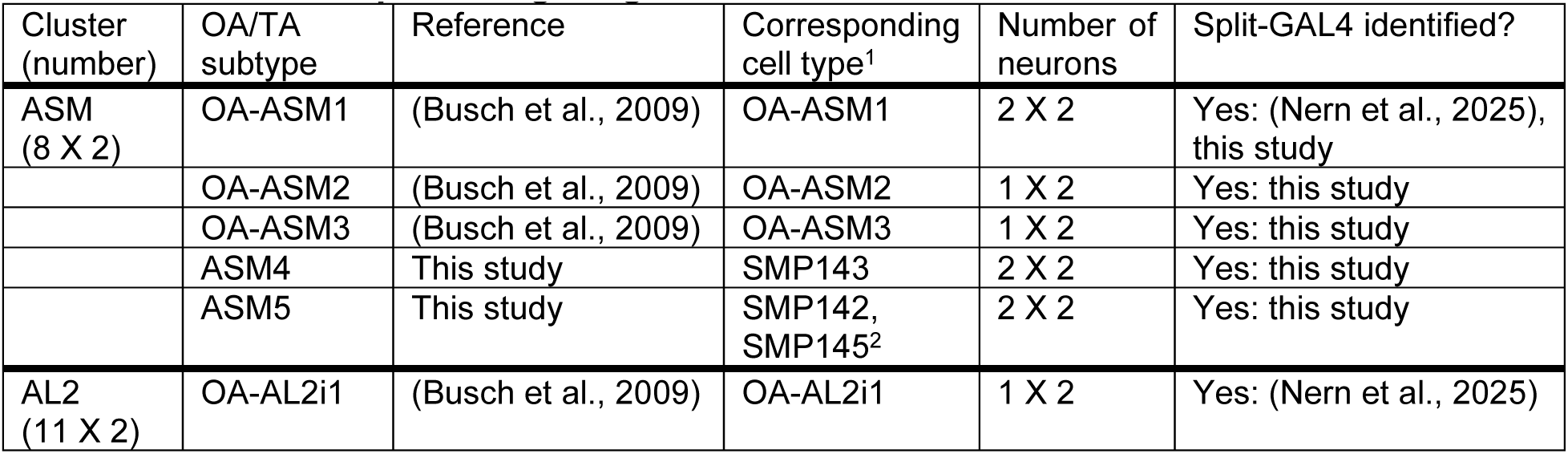
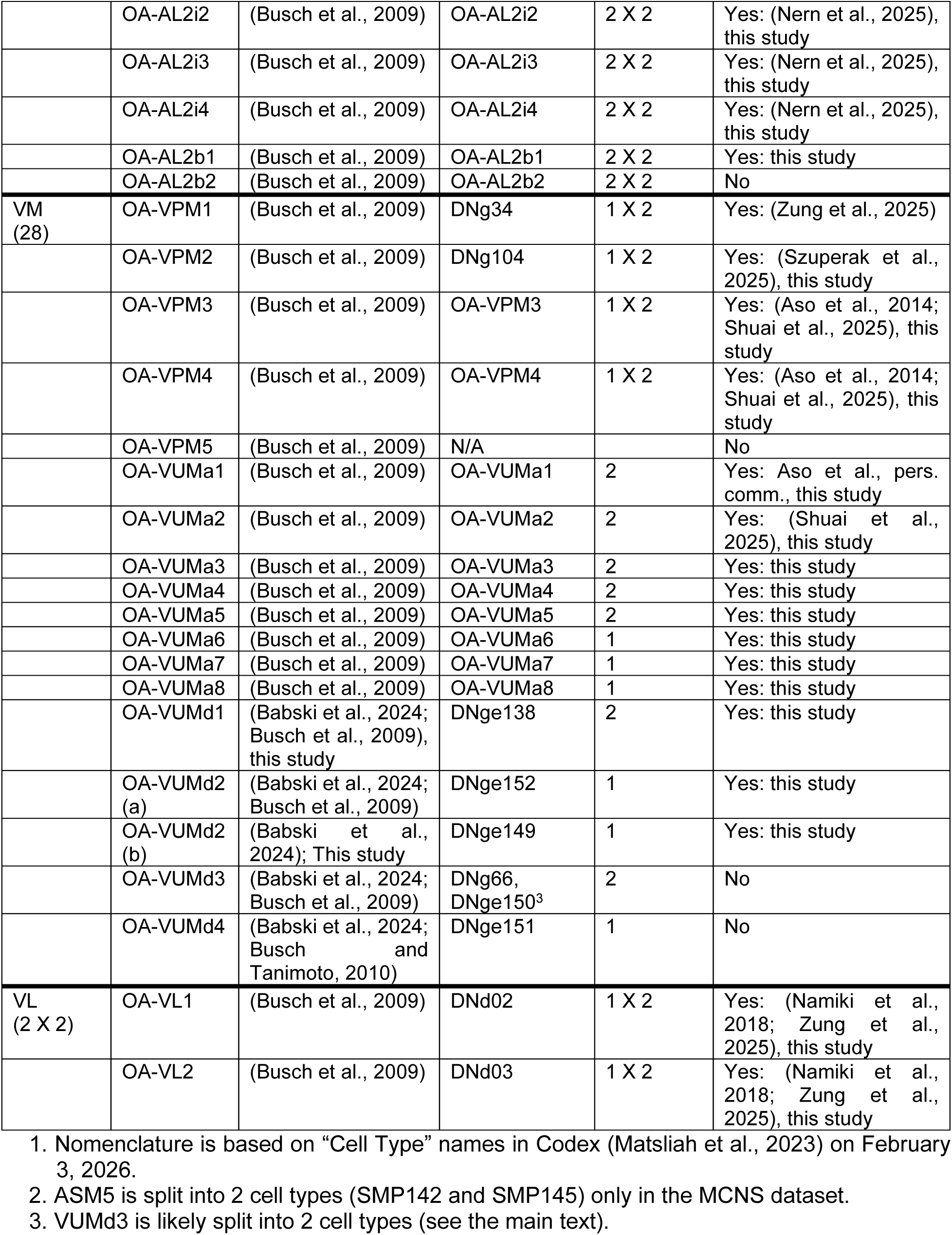
List of *Drosophila* long-range OA/TA neurons.

As detailed below, we observed no gross sex differences in the labeling patterns of the selected split-GAL4 lines. Consistent with this, the number of Tdc2-immunoreactive neurons is similar between males and females, suggesting no obvious numerical sexual dimorphism among OA/TA neurons.

### The ASM cluster contains previously uncharacterized neuronal types

Three distinct neuronal subtypes have been previously identified within the ASM (anterior superior medial) cluster (Busch et al., 2009). ASM1, comprising two neurons per hemisphere, is the only non-AL2 OA/TA neuron that projects to the optic lobe. Its innervation pattern in the optic lobe will be described in the following section along with AL2 neurons. Notably, our ASM1-labeling split-GAL4 lines always co-label neurons in the VL cluster (Fig. 1A, B, Supplementary Fig S2 A, B). By contrast, ASM2 and ASM3 each consist of one neuron per hemisphere, and share highly similar morphology (Fig. 1C, D; Supplementary Fig S2B, C; (Busch et al., 2009)). We were unable to isolate a split-GAL4 line that is specific to either ASM2 or ASM3. Our split-GAL4 lines allowed detailed characterization of ASM neuron innervation patterns, which were only partially described (Busch et al., 2009). The cell bodies of all ASM neurons are located anterior to the superior medial protocerebrum (SMP). From there, ASM1-3 subtypes send their major projections to the posterior side, where extensive innervations are formed across superior, inferior and ventromedial neuropils. ASM1 sends a thick bundle of projections that traverses the posterior neuropils before turning anteriorly to enter the optic lobe. ASM1 also densely innervates a portion of both the superior lateral protocerebrum and SMP. In contrast, ASM2 and ASM3 project ventrally from the posterior side, extensively innervating to the anterior portion of the ventrolateral protocerebrum and to a lesser extent, the ventromedial neuropils. Innervations of ASM1-3 in the central neuropils are biased ipsilaterally, with relatively sparse innervation on the contralateral side.

**Fig. 1.**
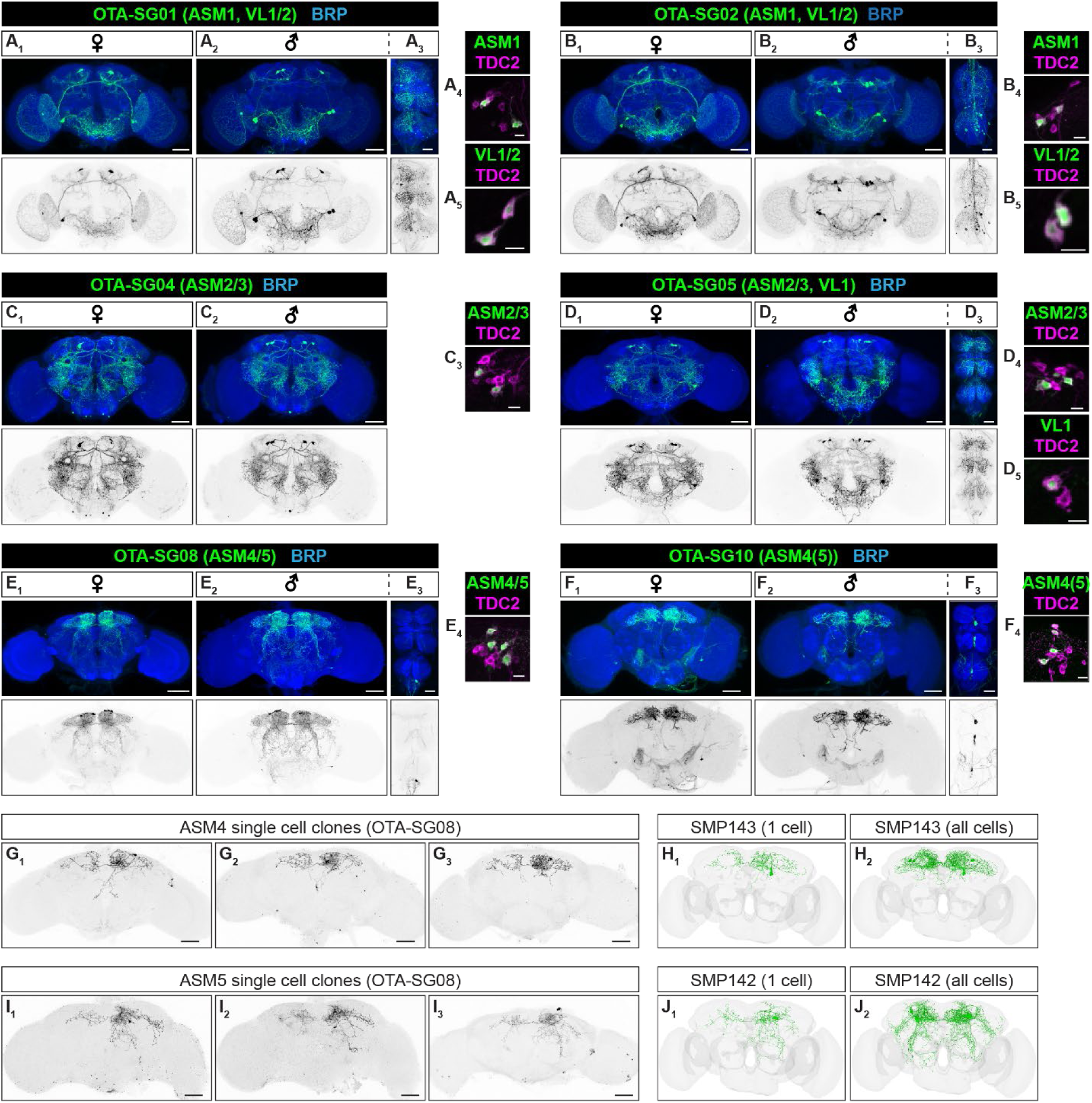
New split-GAL4 reveals two novel ASM subtypes. (**A**-**F**) Representative images of females (**A_1_**, **B_1_**, **C_1_**, **D_1_**, **E_1_**, **F_1_**) and male (**A_2_**, **B_2_**, **C_2_**, **D_2_**, **E_2_**, **F_2_**) brains labeled by ASM neuron-specific split-GAL4 lines (OTA-SG). Images of male ventral nerve cords (**A_3_**, **B_3_**, **D_3_**, **E_3_**, **F_3_**) are also shown for lines that label OA/TA descending neurons (see also Fig. 5). Panels show stacked confocal images, with neuropils and membrane-bound tdTomato expressed by the given split-GAL4 line immunohistochemically visualized by anti-bruchpilot (top panels, blue) and 20XUAS-CsChrimson:tdTomato (top panels, anti-dsRed, green; bottom panels, black and white). Scale bar = 50μm. Targeted cell types are indicated along with a split-GAL4 line name. (**A_4,5_**, **B_4,5_**, **C_3_**, **D_4,5_**, **E_4_**, **F_4_**) Representative images of OA/TA cell clusters targeted by split-GAL4 line, visualized by cytosolic GFP (green), and TDC2 immunoreactivity (magenta). Scale bar = 10μm. (**G**-**L**) Single cell clones from OTA-SG08 obtained with MCFO technique revealed two new ASM types. (**G_1-3_**) Representative images of ASM4 clones, which show extensive innervation of interior region of superior medial protocerebrum (SMP) and superior lateral protocerebrum (SLP). Scale bar = 50μm. ASM4 neurons morphology matches neuronal traces (**H_1_**) and cell number (**H_2_**) of SMP143 cell type in FAFB. (**I_1-3_**) Representative images of ASM5 clones, which innervate the anterior surface of SMP and the posterior lateral protocerebrum, ventrally respect to the lateral horn (LH). Scale bar = 50μm. ASM5 neurons morphology matches neuronal traces (**J_1_**) and cell number (**J_2_**) of SMP142 cell type in the Codex annotation of FAFB.

ASM1-3 subtypes together account for only half of the total ASM neurons (Table 1), which comprise ∼eight Tdc2-expressing cells per hemisphere (Busch et al., 2009; Ishii et al., 2022). Interestingly, we isolated two split-GAL4 lines labelling ∼four *Tdc2*-positive neurons per hemisphere, whose innervation patterns are distinct from any of ASM1-3 neurons (Fig1. E, Supplementary Fig. S2D). Clonal analysis of these lines using MCFO revealed that these four pairs of neurons comprise two distinct subtypes (Fig1. G, I). Strikingly, their morphology resembles neuronal clones isolated from *Tdc2-GAL4*, which encompasses the full OA/TA neurons (Supplemental Fig. S3A; (Chiang et al., 2011; Scheffer et al., 2020)). Corresponding neuronal traces for these two types were identified in the EM volumes, two per hemisphere for each type (Fig. 1H, J; Table 1), consistent with the labeling pattern of our split-GAL4 lines and with the number of missing Tdc2-expressing neurons in the ASM cluster. We named these novel OA/TA cell types ASM4 (SMP143 in Codex annotation) and ASM5 (SMP142/SMP145 in Codex annotation) in continuity with existing nomenclature. Among our drivers, we found another split-GAL4 line that preferentially (though not exclusively) labels ASM4 (Fig. 1F).

Unlike ASM1-3 subtypes, both ASM4 and ASM5 extensively innervate SMP, with clearly distinguishable morphologies (Fig. 1G-J). ASM4 forms dense arborizations within the interior region of SMP and robustly innervates the superior lateral protocerebrum. By contrast, ASM5 arborizes on the anterior surface of SMP, and innervates the posterior lateral protocerebrum just ventral to the lateral horn. ASM5 also projects to a portion of the lateral accessory lobe, which is only weakly innervated by ASM4. Lastly, ASM4 projects to the contralateral hemisphere with a single fiber that crosses the midline above the fan shape body, whereas the midline-crossing fiber of ASM5 runs more anteriorly, above the ellipsoid body.

Neurotransmitter predictions in the Codex database annotate all ASM neurons as dopaminergic; however, Tdc2-expressing ASM neurons do not co-express tyrosine hydroxylase despite their proximity to the PAM dopaminergic neurons (Supplemental Fig. S3B).

### Layer-specific innervation by AL2 neurons suggest precision modulation of visual circuits

Six neuronal subtypes within the AL2 cluster (AL2i1-4 and AL2b1-2) have been characterized in detail (Busch et al., 2009). Briefly, the somata of all subtypes are positioned ventromedially to the antennal lobe and send a single neurite toward the posterior slope where they branch out before projecting to the optic lobe. AL2i1-4 project to the ipsilateral optic lobe while AL2b1-2 send their projections bilaterally (Busch et al., 2009). We isolated split-GAL4 lines that label four subtypes. A previously characterized split-GAL4 driver is used to characterize the neuroanatomy of AL2i1 (Fig. 2A, G, H). Each split-GAL4 line shows relatively high specificity for a single AL2 subtype, although other OA neuronal types (either within AL2 or in the VM cluster) are sometimes co-labeled (Supplementary Fig. S2E, F; see also Supplementary Table S2, S3). A split-GAL4 line for AL2b2 was not identified, although Codex annotates two AL2b2 neurons per hemisphere.

**Fig.2.**
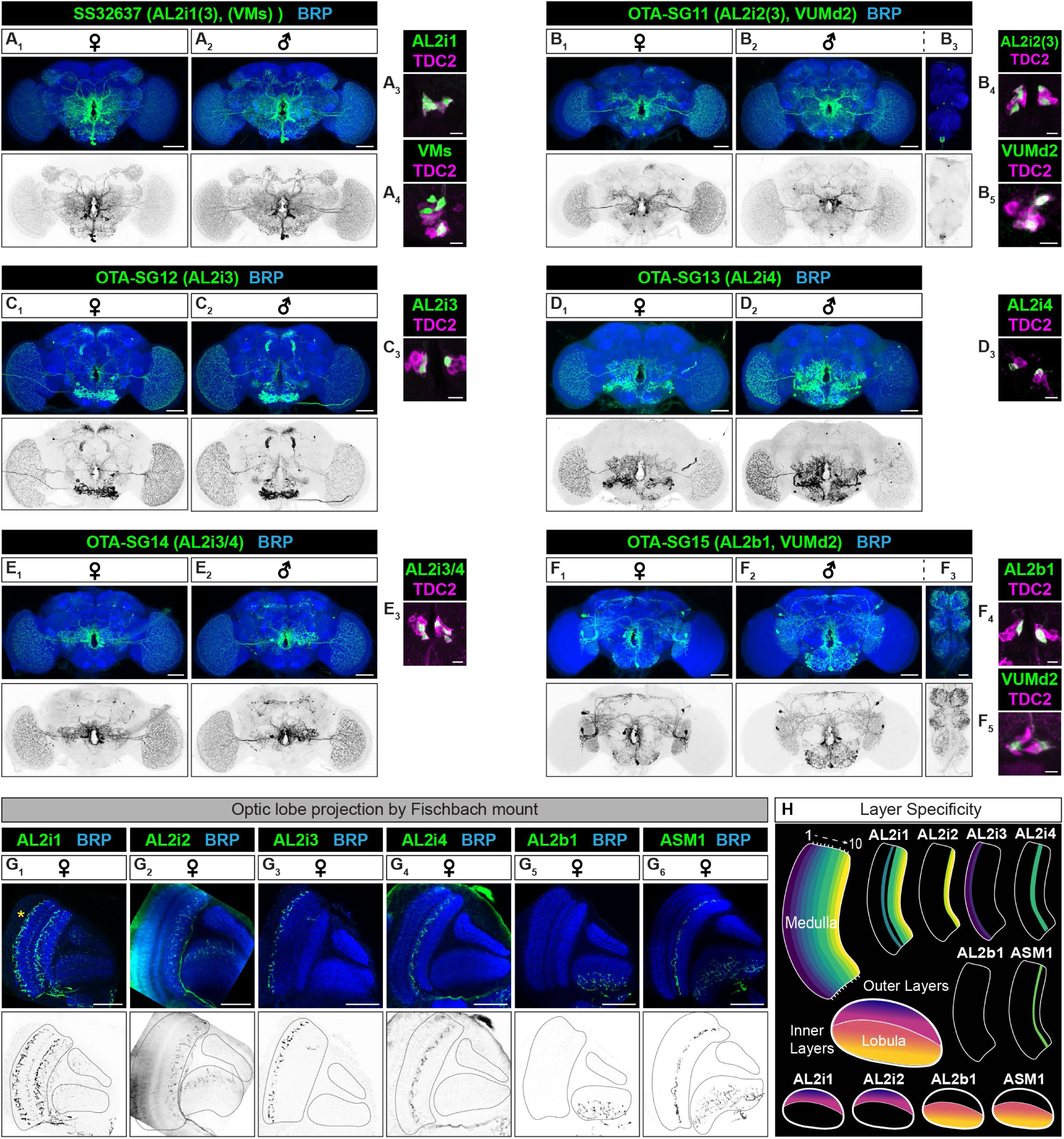
AL2 neurons show layer-specific projections in the optic lobe. (**A**-**F**) Examples of females (**A_1_**, **B_1_**, **C_1_**, **D_1_**, **E_1_**, **F_1_**) and male (**A_2_**, **B_2_**, **C_2_**, **D_2_**, **E_2_**, **F_2_**) brains labeled by AL neuron-specific split-GAL4 lines (OTA-SG). Male ventral nerve cords (**C_3_**, **K_3_**) are also present for lines with OA/TA descending neurons. Panels display stacked confocal images, with neuropils and membrane-bound tdTomato expressed by the given split-GAL4 line shown by immunohistochemical analysis with anti-bruchpilot (top panels, blue) and 20XUAS-CsChrimson:tdTomato (top panels, anti-dsRed, green; bottom panels, black and white) respectively. Scale bar = 50μm. Targeted cell types are indicated along with the split-GAL4 line name. (**A_3,4_**, **B_4,5_**, **C_3_**, **D_3_**, **E_3_**, **F_4,5_**) Representative images of OA/TA cell clusters targeted by split-GAL4 line, visualized by cytosolic GFP (green), and TDC2 immunoreactivity (magenta). Scale bar = 10μm. (**G**) AL2 projection patterns viewed in the dorsal (Fischbach) orientation. Images represent a single z-plane with membrane-bound tdTomato labeling AL2 neurons (green) and Bruchpilot marking synaptic neuropil (blue). In G_1_, Non-AL2i1 innervation of layer 1 is indicated in yellow asterisk (likely AL2i3 neurons). Scale bar = 50μm. (**H**) A schematic summary of the layer-specific projection patterns of AL2 and ASM1 neurons across three neuropils of the optic.

The AL2i1 splitGAL4 driver (SS32637) (Nern et al., 2025) labels three neurons per hemisphere, AL2i1 and likely AL2i3 (Fig. 2A). Notably, this driver also labels multiple Tdc2-immunopositive neurons of the VM cluster (Fig. 2A_4_). Our split-GAL4 driver for AL2i2 (OTA-SG11) labels two neurons per hemisphere, consistent with the connectome (Fig. 2B; (Dorkenwald et al., 2024; Schlegel et al., 2024)). In addition, it stochastically labels one AL2i3 neuron as well as two VUMd2 neurons (Fig. 2B_2_). The AL2i3 split-GAL4 driver (OTA-SG12) labels two AL2i3 neurons per hemisphere (Fig. 2C). We isolated two split-GAL4 drivers for AL2i4 (Fig. 2 D, E). One split driver (OTA-SG13) labels two AL2i4 neurons per hemisphere (Fig. 2D), while the second driver (OTA-SG14) labels 2 neurons per hemisphere for AL2i4 in addition to one AL2i3 neuron expressed stochastically, varying by hemisphere (Fig. 2E). Lastly, our driver for AL2b1 (OTA_SG15) labels two cells per hemisphere as well as VUMd2 neurons (Fig. 2F). Notably, OTA-SG12, SG13, and SG15 also label a moderate number of non-Tdc2-positive cells (Fig. 2C, D, F, Supplementary Table S2, S3).

To further characterize the innervation patterns of AL2 cells in the optic lobe, we imaged brains in the dorsal orientation and annotated the innervation pattern of AL2 neurons as well as optic lobe-projecting ASM1 neurons according to the previously established anatomical framework (Fischbach and Dittrich, 1989). AL2 and ASM1 neurons form a diverse set of layer-specific innervation patterns spanning the medulla, lobula and lobula plate (Fig. 2G-H). AL2i1 neurons exhibited the broadest distribution of all AL2 subtypes, extending processes throughout multiple medulla layers, specifically layers 5, 7, 8, 9, and 10; as well as prominent innervations in the outer lobula and lobula plate (Fig. 2G_1_). AL2i2 neurons showed a more selective innervation profile, with terminals restricted to medulla layers 9 and 10, and outer lobula layers (Fig. 2G_2_). AL2i3 neurons were confined to the most superficial medulla layers 1 and 2 (Fig. 2G_3_) and AL2i4 neurons arborized exclusively within medulla layer 7 (Fig. 2G_4_). AL2b1 neurons are unique in that they lacked medullar projections and innervate only the inner lobula (Fig. 2G_5_). ASM1 neurons innervate layer 8 of the medulla and innermost layers of the lobula (Fig. 2G_6_). Together these findings extend previous anatomical descriptions of AL2 neurons (Busch et al., 2009) by defining their layer specificity within the optic lobe.

### Central brain VM neurons show subtype-specific innervation

Tdc2-expressing cells in the ventral part of the brain consist of two main clusters: ventrolateral (VL) and ventromedial (VM) clusters, classified by the position of their cell bodies within the central brain. Here, we regroup them into 1) neurons that innervate the ventral nerve cord (VNC) (descending neurons), and 2) neurons whose projections are confined to the brain (central neurons). The descending neurons consist of the two VL subtypes, 3-5 VUMd (ventral unpaired medial descending) subtypes, and two VPM (ventral paired medial) subtypes (Babski et al., 2024; Busch et al., 2009), which are described in detail in the following section. Since the remaining neurons all belong to the VM cluster, we call these non-descending neurons central brain VM neurons.

The central brain VM neurons consist of eight VUMa (ventral unpaired medial ascending) subtypes and three of five VPM subtypes. One subtype, however, remains elusive: VPM5 neurons have not been identified in the EM volumes and VPM5 clones have been isolated only three times, despite extensive clonal analyses using *Tdc2-GAL4* (Busch et al., 2009; Busch and Tanimoto, 2010; Chiang et al., 2011). We were unable to identify a split-GAL4 line that labels this class.

Interestingly, most split-GAL4 lines selective for central VM neurons label multiple subtypes, likely reflecting their shared neurodevelopmental origin (Busch and Tanimoto, 2010). Our split-GAL4 lines complement previously reported GAL4 lines that label specific subtypes in this cluster (Aso et al., 2014; Shuai et al., 2025), collectively allowing us to examine cell-type specific functions of all central VM neurons in a combinatorial manner.

Central brain VM neurons are arguably among the most extensively innervating neurons in the fly brain (Busch et al., 2009), although there are qualitative differences among them. VUMa1-5 (Fig. 3A-E, Supplementary Fig. S2E-K) show innervations to relatively restricted and well-defined neuropils. These neurons exist as doublets, but their differences are detectable only in the position of soma, reflecting their distinct lineages (Busch and Tanimoto, 2010). In contrast, VUMa6-8 (Fig. 3A, F-H, Supplementary Fig. S2K, L, N), as well as VPM3 and VPM4 (Fig. 3H-J, Supplementary Fig. S2L, N, O), innervate multiple brain regions and neuropils often via diffuse fibers. In particular, VUMa6-8 widely innervate neuropils in the posterior side of the brain, most of which have not been functionally well-characterized (Scheffer et al., 2020). Their gross innervation areas in this region overlap even though their neuronal morphologies are distinct.

**Figure 3:**
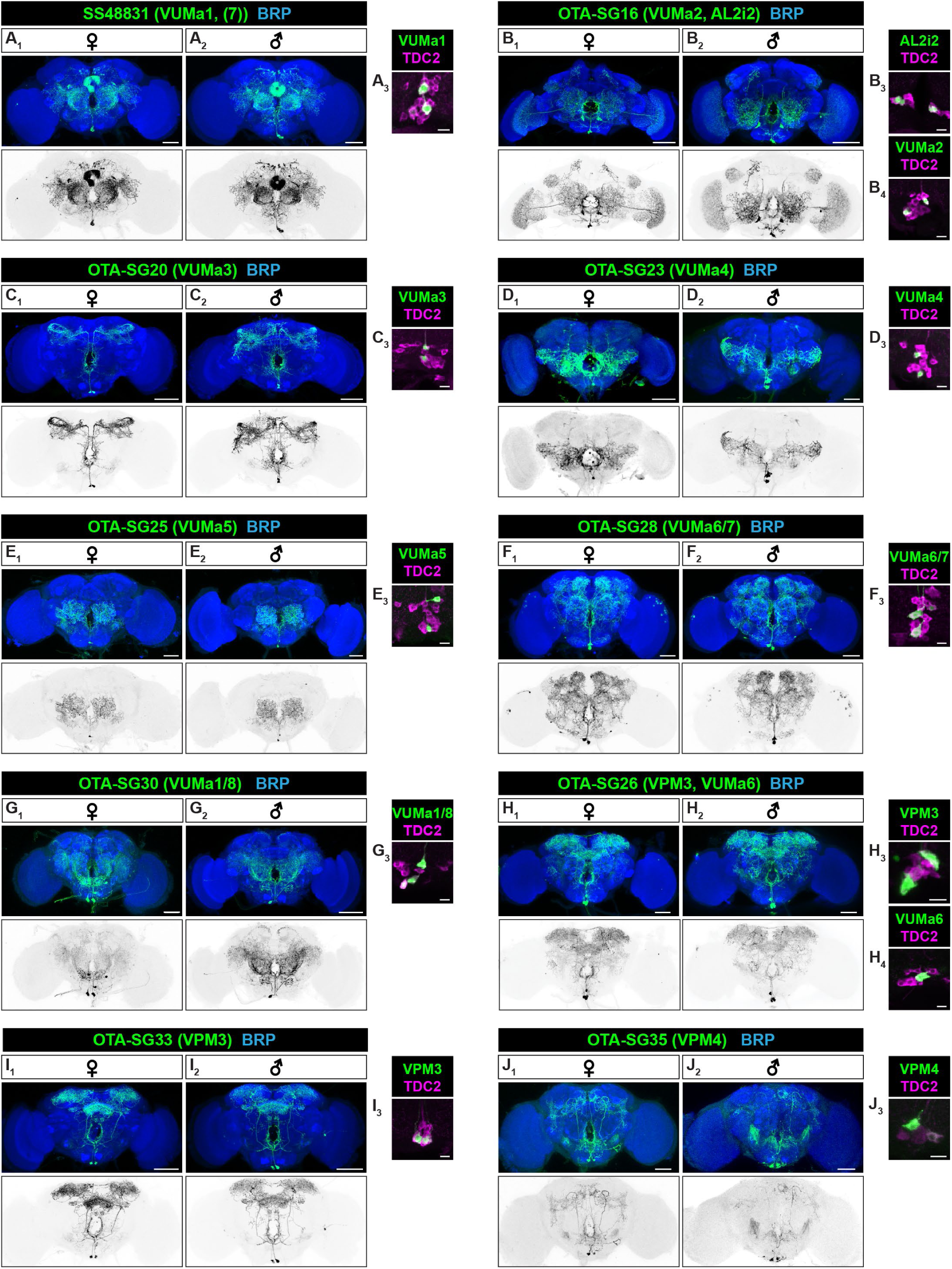
New split-GAL4 lines for combinatorial targeting of VUMa and VPM3/4 neurons. Immunohistochemical analysis of females (**A_1_-J_1_**) and male (**A_2_-J_2_**) brains labeled by split-GAL4 lines (OTA-SG) with high specificity for VUMa1-8, VPM3, and VPM4 cell types. Targeted cell types are indicated along with the split-GAL4 line name. Panels show maximum intensity projections, with neuropils and membrane-bound tdTomato expressed by the given split-GAL4 line visualized by anti-bruchpilot (top panels, blue) and 20XUAS-CsChrimson:tdTomato (top panels, anti-dsRed, green; bottom panels, black and white) respectively. Scale bar = 50μm. (**A_3_**, **B_3,4_**, **C_3_**, **D_3_**, **E_3_**, **F_3_, G_3_, H_3,4_, I_3_, J_3_**) Representative images of OA/TA cell clusters targeted by split-GAL4 line, visualized by cytosolic GFP (green), and TDC2 immunoreactivity (magenta). Scale bar = 10μm.

### Heterogeneity in OA/TA descending neurons

Descending neurons represent a crucial bottleneck, linking sensory pathways with motor circuits (Namiki et al., 2018). As briefly described above, at least seven wide-projecting OA/TA subtypes are descending neurons. Among these, we confirmed that previously characterized split-GAL4 lines labeling VL (ventrolateral)-like neurons ((Namiki et al., 2018), Fig. 4A) indeed express Tdc2 (Fig. 4A_4_). We also identified other split-GAL4 lines that target both VL1 and VL2 subtypes (Fig. 4B, Supplementary Fig. S2A, B).

**Figure 4:**
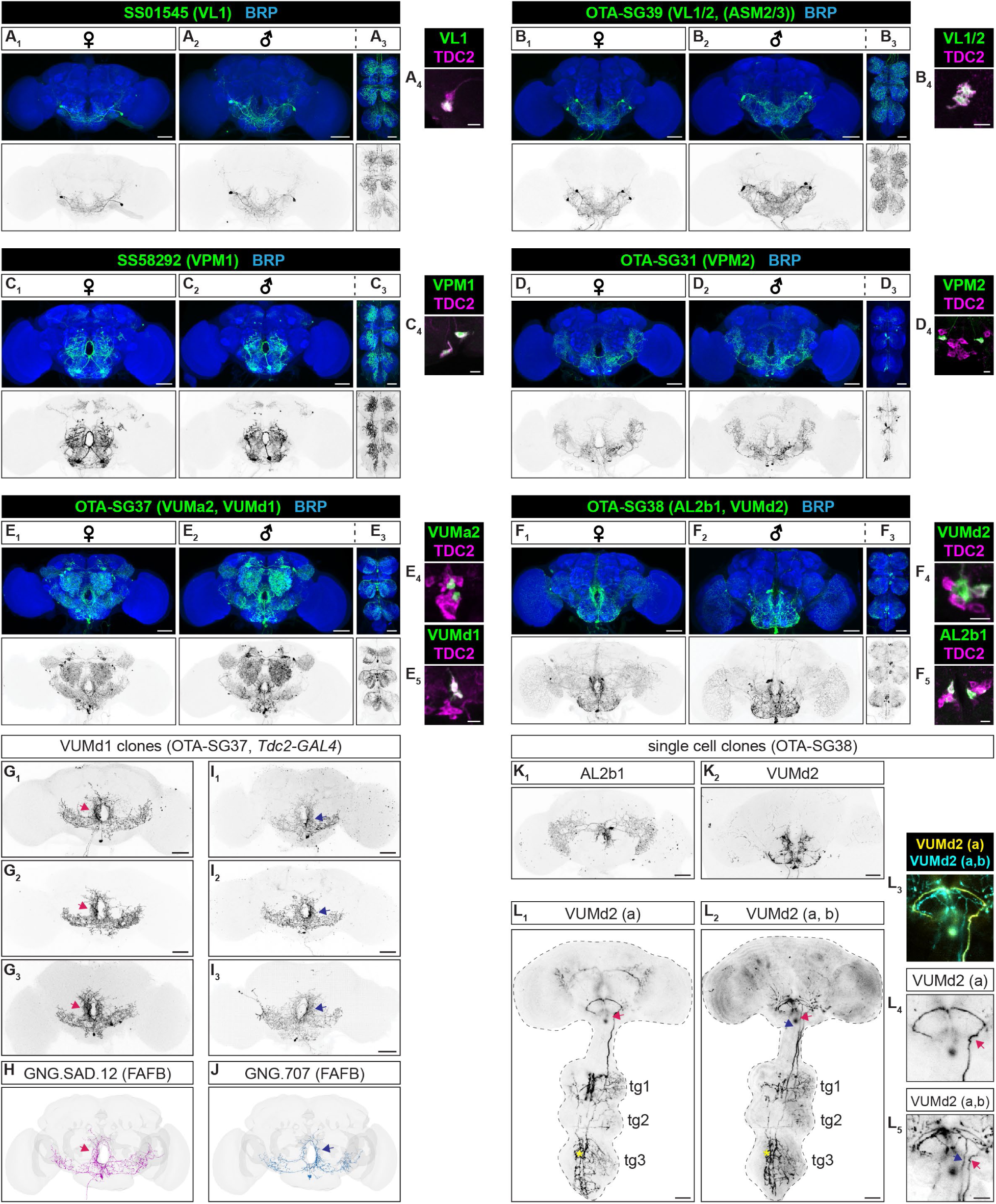
Neuroanatomical diversity of descending OA/TA neurons. (**A**-**F**) Representative images of females (**A_1_**, **B_1_**, **C_1_**, **D_1,_ E_1_**, **F_1_**) and male (**A_2_**, **B_2_**, **C_2_**, **D_2,_ E_2_**, **F_2_**) brains labeled by descending OA/TA neuron-specific split-GAL4 lines. Images of male ventral nerve cords (**A_3_**, **B_3_**, **C_3_**, **D_3,_ E_3_**, **F_3_**) are also shown. Panels show maximum intensity projections, with neuropils and membrane-bound tdTomato expressed by the given split-GAL4 line visualized by immunohistochemical analysis with anti-bruchpilot (top panels, blue) and 20XUAS-CsChrimson:tdTomato (top panels, anti-dsRed, green; bottom panels, black and white) respectively. Scale bar = 50 μm. Targeted cell types are indicated along with the split-GAL4 line name. (**A_4_**, **B_4_**, **C_4_**, **D_4,_ E_4,5_**, **F_4,5_**) Representative images of OA/TA cell neurons targeted by split-GAL4 line, visualized by cytosolic GFP (green), and TDC2 staining (magenta). Scale bar = 10 μm. (**G-J**) VUMd1 shows asymmetric innervations around the esophagus. (**G**) Representative images of single cell clones of VUMd1 neurons with thicker innervations on the left side obtained from OTA-SG37 (**G_1_**) and from Tdc2-GAL4 (**G**_2,3_) with MCFO technique in male flies, corresponding to GNG.SAD.12 in FAFB (**H**). (**I**) Clones of VUMd1 neurons with right bias innervations, obtained from Tdc2-GAL4 (**I**_1,2_) and OTA-SG37 (**I_3_**) with MCFO technique in male flies, corresponding to GNG.707 in FAFB (**J**). Scale bar = 50 μm. (**K**) Representative image of single cell clones of AL2b1 neurons (**K**_1_) and VUMd2 (**K**_2_) obtained from OTA-SG38 with MCFO technique in male flies. Scale bar = 50 μm. (**L**) Clonal analysis suggests the existence of two distinct VUMd2 subtypes. VUMd2 (a) (**L_1_**) present one lateral descending projection each side (only one side was visible in this sample) from the lateral inferior posterior slope. VUMd2 (b) (**L**_2-5_) presents additional finer neurites that run more centrally in the cervical connective (arrow, L_4,5_). Non-VUMd2 neurons are indicated in yellow asterisks. Scale bar = 50 μm (L_1,2_), 10 μm (L_5_).

OA/TA neurons that express a sex-determining gene *fruitless* (*fru*) (Andrews et al., 2014; Certel et al., 2010) are only found among OA/TA descending neurons (VPM1, VUMd3). We found four FruM+ Tdc2-expressing neurons in the VM cluster (Supplemental Fig. S4A), which is consistent with the number of VPM1 and VUMd3 neurons annotated in Codex. While VPM1-labeling split-GAL4 lines have been described elsewhere (Fig. 4C; (Zung et al., 2025)), no genetic driver that specifically labels the VUMd3 subtype has yet been documented. The VPM2 subtype, recently recognized as a descending neuron (Babski et al., 2024) consists of a pair of neurons with ipsilateral brain projections and a single neurite innervating the VNC (Fig. 4D, Supplementary Fig. S2M) (Szuperak et al., 2025).

Detailed characterization of split-GAL4 lines labelling VUMd1 and VUMd2 revealed neuroanatomical complexity that has been inferred from stochastic labeling with dye injection (Babski et al., 2024). Although VUMd1 was originally described as an “unpaired” (bilaterally symmetrical) neuron, our MCFO-generated clones of VUMd1 from both the OTA-SG37 split-GAL4 line and *Tdc2-GAL4* show reproducible asymmetry in the innervations to the suboesophageal region (Fig. 4G, I). Each VUMd1 neuron sends the thicker projection to one side of the esophagus, resulting in denser innervation on that side. VUMd1 neurons correspond to DNge138 in Codex, and are present as a bilateral doublet in FAFB and MCNS datasets. Notably, in both EM volumes, one DNge138 innervates more densely on the right side of the brain, while the other neuron shows the opposite innervation bias (Fig. 4H, J).

Regarding VUMd2, we found evidence that this OA/TA subtype can be subdivided into two neuroanatomically distinct neurons, consistent with a previous observation (Babski et al., 2024). VUMd2 was originally reported to have two descending projections entering the cervical connective laterally from the lateral inferior posterior slope (Busch et al., 2009). DNge149, one of the two neurons annotated as VUMd2 in Codex (VUMd2 #1 in (Babski et al., 2024)), matches this morphology (Supplementary Fig. S4B-D). In contrast, another VUMd2 cell named DNge152 (VUMd2 #2 in (Babski et al., 2024)) shows additional fine neurites running posteriorly along the midline of the cervical connective, in both BANC (female) and MCNS (male) volumes (Supplementary Fig S4E-G). Consistent with these findings, our clonal analysis of the VUMd2-labeling OTA-SG38 line (Fig. 4F, K), which labels two Tdc2-expressing neurons in the VM cluster (Fig. 4F_4_), revealed two distinct subtypes, with descending projections similar to those described in the connectome datasets (Fig. 4L).

Heterogeneity in descending projections was also observed in VUMd1 neurons: one clone type had two parallel descending projections with non-overlapping target neuropils in the VNC (Supplementary Fig. S4H), while the other type had only one descending projection that exits the brain (Supplementary Fig. S4I). DNge138 in FAFB (Fig. 4H, J) and BANC (Supplementary Fig. S4J, K) contains these two types, but both DNge138 neurons in MCNS have two parallel projections (Supplementary Fig. S4L, M). It is unclear whether the differing number of descending projections reflect distinct functional subtypes within VUMd1 neurons, or it is a result of developmental stochasticity. The number of descending projections does not correlate with the laterality of innervations proximal to the esophagus.

Two VUMd3-like neurons have been described previously (Babski et al., 2024; Szuperak et al., 2025), and one VUMd4 neuron have been reported (Babski et al., 2024; Busch and Tanimoto, 2010). These two cell types have not been extensively characterized due to the lack of cell type-specific genetic drivers (Babski et al., 2024; Busch and Tanimoto, 2010; Szuperak et al., 2025). Because we did not succeed in isolating split-GAL4 lines for these cell types, detailed characterization of VUMd3 and VUMd4 subtypes will require further studies.

### OA/TA neuron cell types have distinct connectivity similarity

The two new cell types in the ASM cluster and the heterogeneity within the VUMd subclass described above prompted us to re-examine OA/TA cell types based on the synaptic connectivity revealed by the EM volumes. To this end, we calculated the connectivity similarity between each of 62 candidate long-range OA/TA neurons in female (FAFB) and male (MCNS) EM volumes based on cell type-based synapse counts (see Methods for details).

In general, cosine similarity was higher between neuronal traces belonging to the same cell type than between cells from different cell types, forming characteristic series of “squares” with higher values along the diagonal line of the matrix in both female (Fig. 5A) and male (Fig. 5B) datasets. The similarities within each cell type calculated using downstream cell types are qualitatively higher than values using upstream cell types (Fig. S5A, B). There are more downstream cell types than total upstream cell types, which may reflect OA/TA neurons’ role to sort overlapping inputs to discrete outputs. Because similar connectivity suggests similar function in the circuit, this observation reinforces the notion that neuroanatomically defined OA/TA cell types represent distinct functional units in the fly brain.

**Figure 5:**
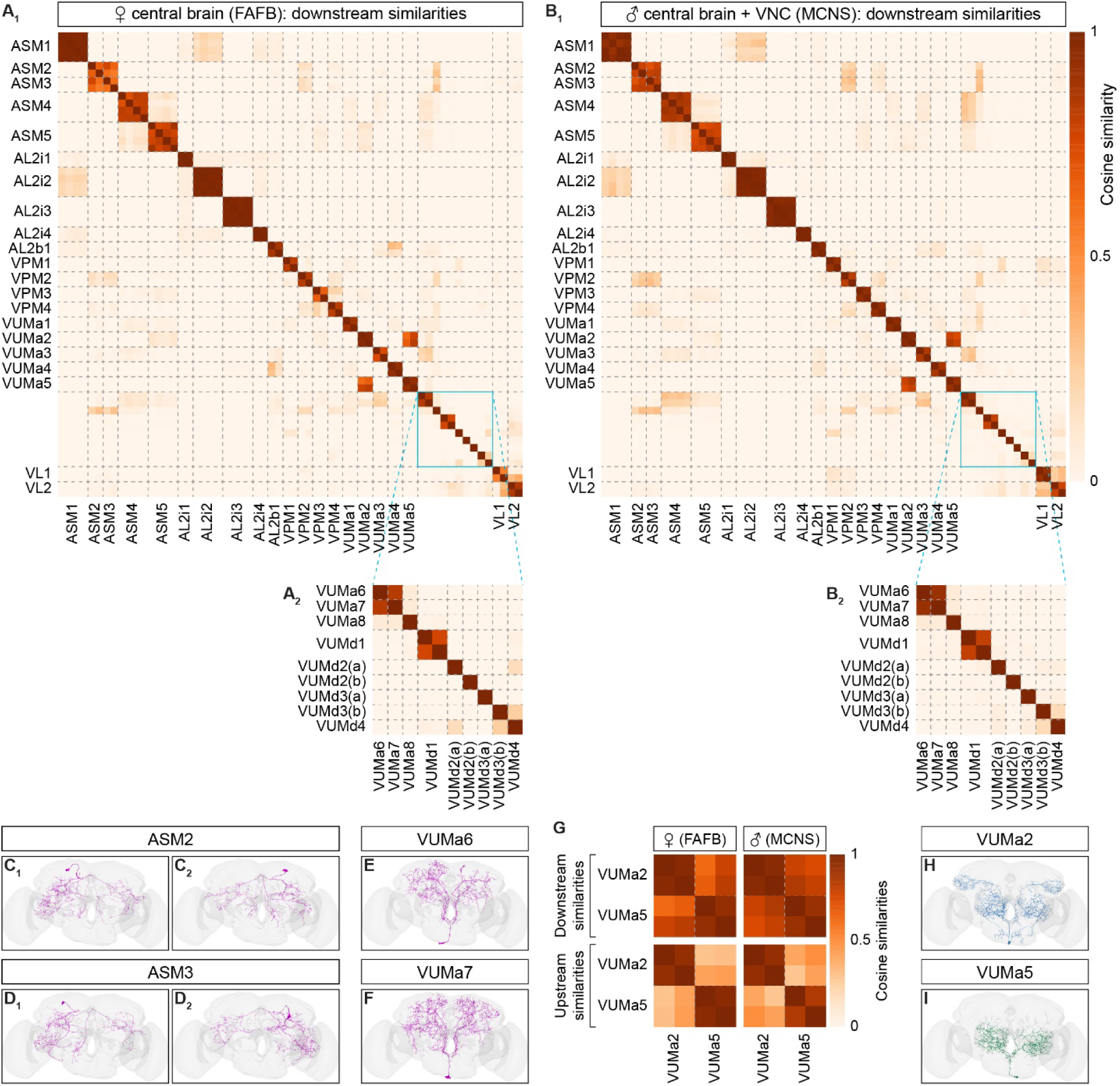
Connectivity similarity reflects OA/TA cell types. (**A, B**) Connectivity similarity of OA/TA neurons identified in the EM volumes of the female central brain (FAFB) (**A**) and the male central brain and nerve cord (MCNS) (**B**) based on their downstream neuronal types. Broken lines represent the cell type boundaries. Regions that include a cell type containing only one neuron (cyan squares in **A_1_**, **B_1_**) are enlarged below (**A_2_**, **B_2_**). (**C-E**) Images of neuronal traces of ASM2 (**C**), ASM3 (**D**), VUMa6 (**E**), and VUMa7 (**F**) neurons in FAFB. **G**. Connectivity similarities of VUMa2 and VUMa5 neurons in FAFB (left) and MCNS (right) EM volumes, based on their downstream (top) and upstream (bottom) neuronal types. (**H, I**) Images of neuronal traces of VUMa2 (**H**) and VUMa5 (**I**) neurons in FAFB.

However, we found several notable exceptions. First, the connectivity between ASM2 and ASM3, and between VUMa6 and VUMa7, are highly similar (Fig. 5A, B, Supplementary Fig. S5A, B), which parallels their morphological similarities (Fig. 5C-F). The similarity levels are comparable to those observed within other cell types, suggesting that these neurons form single computational units from a functional perspective. We also noticed that VUMa2 and VUMa5 show a high downstream connectivity similarity (Fig. 5A_1_, B_1_), likely because they share similar downstream neurons in the antennal lobe. However, relatively low upstream similarities between these two subtypes distinguish them (Fig. 5G), consistent with their overall neuroanatomical differences (Fig. 5H, I). Namely, VUMa2 has a distinct bilateral projection to both mushroom bodies and lateral horns, which is absent in VUMa5 (Busch et al., 2009).

The two VUMd1 neurons (DNge138) show high connectivity similarity (Fig. 5A_2_, B_2_, Supplementary Fig. S5A_2_, B_2_,), which aligns with our neuroanatomical analysis suggesting that this class exists as a pair of neurons. In contrast, neuronal volumes of the two VUMd2, two VUMd3 candidates and one VUMd4 candidate all show distinct connectivity patterns (Fig. 5A_2_, B_2_, Supplementary Fig. S5A_2_, B_2_,), suggesting that each represents a functionally unique cell type, as previously suggested (Babski et al., 2024; Szuperak et al., 2025). Because these classes of descending neurons make relatively few synapses in the central brain (Babski et al., 2024), we also created the similarity matrix using the MANC (male nerve cord) EM volume (Takemura et al., 2023), which only contains synapses in the male VNC. Consistent with the results from FAFB and MCNS, neural volumes assigned to VUMd1 show high connectivity similarities while volumes assigned to two VUMd2 neurons, two candidate VUMd3 neurons, and one candidate VUMd4 neuron are all distinct from one another (Supplementary Fig. S5C, D). Overall, our connectome data analysis prompts us to adjust a few previously proposed OA/TA cell type classifications based on their circuit functions.

### Short-range OA/TA neurons

As noted above, Tdc2 is also expressed in neurons outside the set of four long-range OA/TA clusters. The Tdc2 immunoreactivity of a putative AL1 cluster (one cell per hemisphere; (Busch et al., 2009)) was unreliable and weak in our preparations. Aside from the AL1 cluster, cell bodies of Tdc2 positive neurons are predominantly located on the posterior side of the brain. The identity of this posterior cluster difficult to access as we were unable to generate single-cell clones using *Tdc2-GAL4,* possibly due to the relatively weak labeling of these neurons by this driver.

We detected consistent co-labeling of Tdc2 and GFP driven by *Vmat^GAL4^* in all of the long-range OA/TA neuronal clusters as previously mentioned (Supplementary Fig. S1A), as well as in many neurons with small cell bodies in the posterior medial part of the brain (Supplementary Fig. S6A). These neurons likely correspond to the PB1 cluster (Busch et al., 2009). Aside from PB1 cells, we did not find other small Tdc2-immunoreactive cells that are also co-labeled by *Vmat^GAL4^*. These include PSM Supplementary Fig. S6B) and PB2 cells (Supplementary Fig. S6C), which are neurons with large somata that were previously reported to be immunoreactive to anti-octopamine antiserum (Busch et al., 2009). Thus, the PB1 cluster may be the only OA/TA-releasing neurons that do not belong to the long-range subtypes described thus far. These PB1 neurons likely correspond to eight fan shape body-projecting OA/TA neurons (Wolff et al., 2025), which are 73 in total according to the FAFB annotation. These neuronal subtypes project relatively locally inside the fan shape body, in contrast to the long-range projection of OA/TA neurons (Wolff et al., 2025). Neuronal clones with morphologies similar to these “short-range” cell types were previously isolated from *Tdc2-GAL4* (Chiang et al., 2011). Aside from the EL subtype, none of these short-range, fan shape body-projecting OA/TA neurons express *Tbh* (Wolff et al., 2025), suggesting that they are likely tyraminergic.

### ASM4/5 neurons acutely suppress aggression

OA/TA neurons can be divided into two subsets with opposing effects on *Drosophila* aggression. While enhanced signaling in the entire OA/TA neurons (*Tdc2-GAL4*) was reported to increase aggression (Zhou et al., 2008), the activation of *Tdc2-GAL4* neurons co-expressing the transcriptional regulator *nervy* (*nvy*) suppresses aggression among socially isolated flies (Ishii et al., 2022). The *nvy*-expressing OA/TA neurons mostly (but not exclusively) belong to the ASM and VL clusters (Ishii et al., 2022). Using the newly established genetic access to OA/TA subtypes, we sought to address whether this aggression-suppressing function could be attributed to a specific OA/TA subtype.

We expressed a red-shifted channelrhodopsin CsChrimson (Klapoetke et al., 2014) using split-GAL4 lines targeting neurons in the ASM and VL clusters. Following social isolation, targeted neurons were stimulated with intermittent 655nm LED light, and their level of aggression were quantified (Fig. 6A). Optogenetic activation of VL neurons and ASM1-3 neurons did not decrease aggression relative to genetic controls (Fig. 6B). In striking contrast, for the flies that express CsChrimson in ASM4/5 neurons, fighting was acutely and reversibly suppressed during LED stimulation period compared to both the post-stimulation period and genetic controls during stimulation period. Suppression by the activation with the OTA-SG10 line, which preferentially labels ASM4 neurons, suggests that ASM4 may be more important for aggressive behavior. Importantly, this phenotype was observed in both male (Fig. 6B) and female (Fig. 6C) flies. Immunoreactivity with Nvy protein was detectable in all ASM and VL neurons (Fig. 6D), consistent with our previous observation (Ishii et al., 2022). Our novel GAL4 lines enabled us to identify the OA/TA subtypes that regulate aggressive interactions among flies.

**Figure 6:**
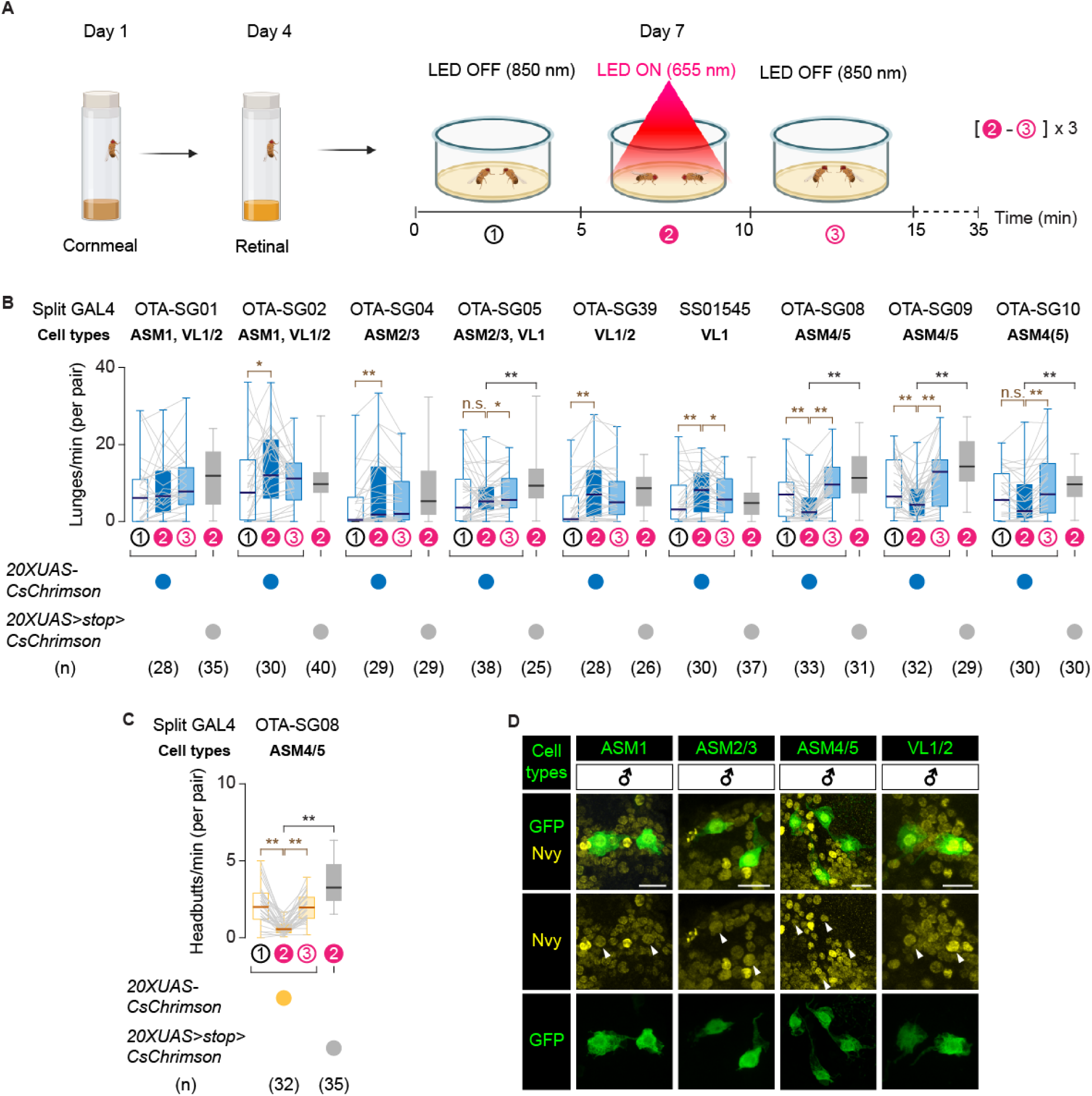
ASM4/5 neurons acutely suppress aggression among socially isolated flies. (**A**) Schematic representation of optogenetic assay to measure aggressive behavior in socially isolated flies (6 days isolation), in the indicated time period. Pre-stimulation (1), optogenetic activation (2), post-stimulation (3). (**B-C**) Boxplot of lunges/minutes (males, **B**) and headbutts/minutes (females, **C**) by the tester flies expressing either 20XUAS-CsChrimson:tdTomato or 20XUAS>stop>UAS-CsChrimson:tdTomato (negative control) by the indicated split-GAL4. Optogenetic stimulation of ASM and VL -OA subtypes show reduced aggression only in ASM4/5 lines compared with prestimulation period and with negative controls during stimulation. Tester genotypes and pairs are indicated below the plot. Experimental flies pre(1)- ON(2)- OFF(3) were compared using Wilcoxon’s signed rank test (with Bonferroni correction). Experimental flies and negative controls (UAS>stop>CsChrimson, during ON period), were compared using Mann-Whitney U test. ns *p > 0.05, * p < 0.05, ** p < 0.01. (**D**) Representative images of indicated OA/TA cell types, visualized by cytosolic GFP (green), and *nvy* immunoreactivity (yellow). Scale bar = 10 μm.

### AL2i1 and AL2i2 differentially modulate visual fight control

We tested two common visual behaviors during flight - optomotor reflexes that reduce retinal slip to stabilize the direction of gaze, and object orientation responses that guide the animal toward salient visual features - while activating or silencing AL2i1(SS32637) and AL2i2 (OTA-SG11) OA/TA subtypes. Flies were tested in a virtual reality flight simulator (Fig. 7A). Revolving freely on a magnetic pivot, the animal has closed-loop control over its visual and proprioceptive dynamics. To test the optomotor reflex, we used a rotating 360-degree background moving at constant velocity (Fig. 7A). To test object orientation, we used a 30° bar composed of the same random texture rotating across a stationary background.

**Figure 7:**
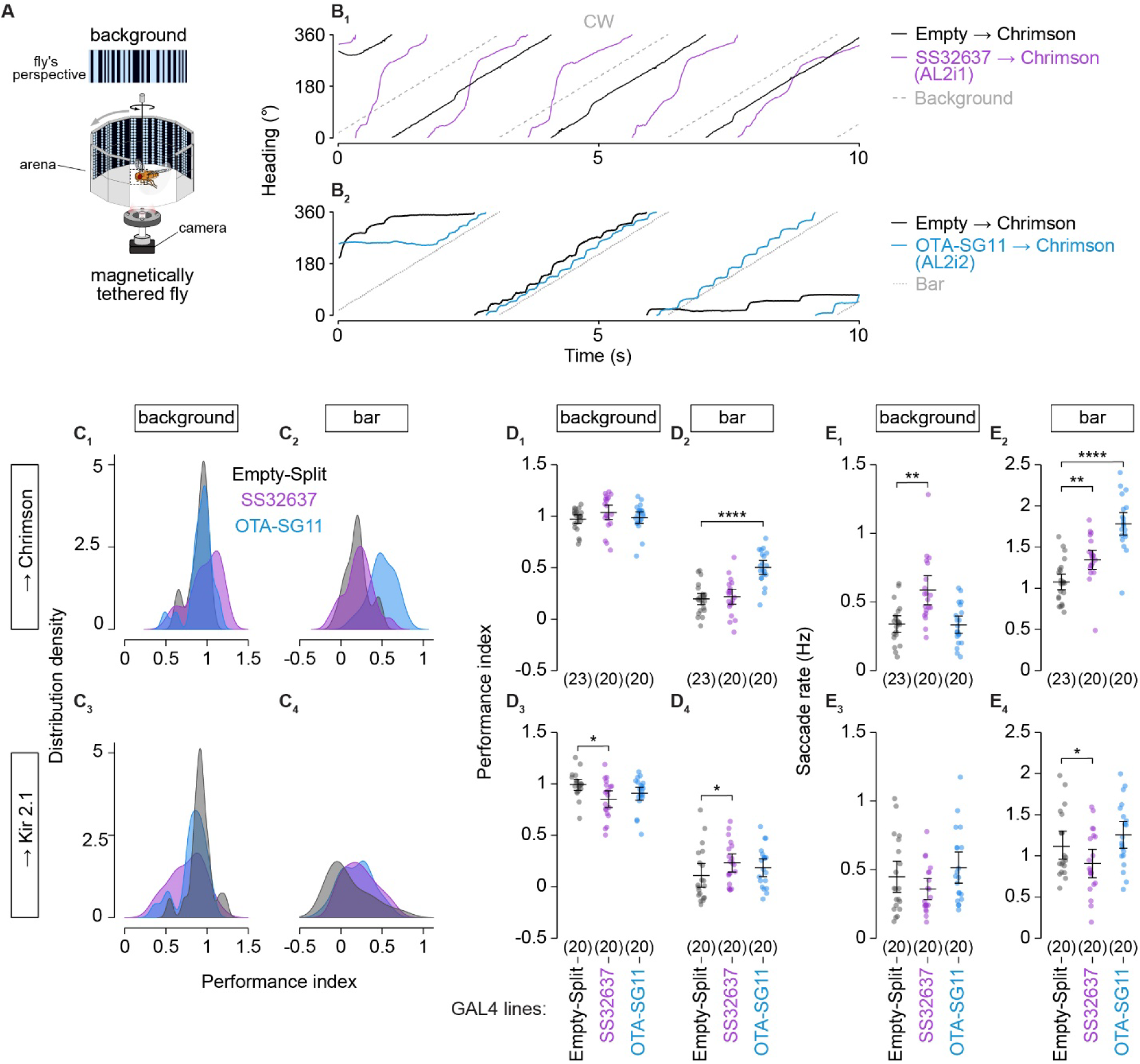
AL2i1 and AL2i2 differentially modulate visual flight control. **(A)** Top: A representation of the fly’s visual perspective, a randomized pattern of bright and dark stripes. Bottom: A schematic of the magnetic-tether setup where a fly is attached to a stainless-steel pin using UV-curable glue and suspended within a magnetic field at the center of a 360° LED arena. The magnetic tether allows unrestricted rotation about the fly’s yaw axis. Infrared illumination is provided from below, and a bottom-mounted camera records the fly’s behavior. **(B)** Wrapped raw heading traces plotted against stimuli position (dashed grey) for flies expressing CsChrimson either in SS32637 (AL2i1, B_1_, purple traces) or OTA-SG11 (AL2i2, B_2_, blue traces). **(C)** Performance index (PI) is defined as the ratio of the fly’s angular displacement to that of the visual stimulus at the end of one trial. P = 1 indicates perfect tracking of the stimulus, PI < 1 reflects reduced tracking performance. PI > 1 when flies overshoot the stimulus. Distribution density curves of PI values are plotted for flies in six different genotypic groups presented with either a revolving randomized background (C_1_ and C_3_) or a revolving bar (C_2_ and C_4_). Genotypes include: a promoter-less driver, SS32637(AL2i1), and OTA-SG11(AL2i2) driving either CsChrimson (C_1_ and C_2_) or Kir2.1(C_3_ and C_4_). **(D)** Scatter plots of mean PI over three repetitions per fly for clockwise (CW) and counterclockwise (CCW) trials for six genotypes under two visual conditions. **(E)** Scatter plots of mean frequency over three repetitions per fly for clockwise (CW) and counterclockwise (CCW) trials for all genotypes under two visual conditions. For D and E, black bars indicate the mean and the bootstrapped 95% confidence interval. Sample numbers are indicated in parenthesis. * p < 0.05, ** p < 0.01, **** p < 0.0001 by unpaired t-test after Bonferroni corrections.

Single fly traces show that control flies tend to steer smoothly, at constant body velocity, while also occasionally making rapid body turns (saccades) to stabilize panoramic background motion (Fig. 7B_1_). By contrast, control flies track the bar only with saccades when the bar is near the frontal field of view, and remain fixated on the stationary visual background between saccades (Fig. 7B_2_). Optogenetic activation of AL2i1 (SS32637) neurons with CsChrimson caused a fly to “over stabilize” its gaze by rotating faster than the stimulus (Fig. 7B_1_). By contrast, optogenetic activation of AL2i2 (OTA-SG11) neurons elicited more body saccades, which strengthens bar tracking (Fig. 7B_2_).

To quantify the influence of depolarizing or hyperpolarizing OA/TA subtypes on these visual flight behaviors, we computed a performance index (PI) and measured saccade rate (see Methods for details). The only manipulation that influenced PI strongly was the optogenetic activation of AL2i2 (OTA-SG11), which significantly increased bar tracking performance by shifting PI closer to 1 (Fig. 7C_2_, D_2_).

Optogenetic depolarization of neither AL2i1 nor AL2i2 by CsChrimson had any significant effect on smooth optomotor responses to a background panorama (Figure 7C_1_, D_1_) – the average responses were already essentially saturated (PI=1); it is worth noting that high optomotor gain is typical for flies. Activating AL2i1 increased saccade rates during both wide-field background and bar motion experiments (Fig. 7C_1_, C_2_, E_1_, E_2_). An increase in saccade rate without concomitant increased tracking performance reflects more undirected saccades. By contrast, optogenetic activation of AL2i2 resulted in robustly increased bar tracking performance (Fig. 7C_2_, D_2_) accompanied by a very strong increase in saccade rate (Figure 7E_2_) indicating more bar-directed, tracking saccades when this OA/TA line is photoactivated.

Constitutive hyperpolarization of AL2i1 by Kir2.1 resulted in a modest reduction of background optomotor responses (Fig. 7C_3_, D_3_), without a significant accompanying change in saccade rate (Figure 7E_3_). Silencing AL2i2 modestly increased bar tracking performance (Fig. 7C_4_, D_4_) with fewer saccades (Fig. 7E_4_) suggesting slightly higher smooth optomotor tracking with fewer object tracking saccades. Although SS32673 and OTA-GS11 do not selectively label AL2i1 and AL2i2, respectively, AL2i3 is the only additional AL2 subtype labeled by both drivers. Thus, the distinct effects observed are unlikely to be explained by manipulation of AL2i3 and instead point to contributions from AL2i1 and AL2i2.

## Discussion

OA has been described to prompt the organism into “dynamic action” (Verlinden et al., 2010). A growing body of evidence shows that OA/TA neurons are crucial for the modulation of multiple processes in invertebrates. Taking advantage of our newly established, comprehensive genetic tool, we demonstrate that the role of OA/TA modulation on a specific behavioral process can be indeed attributed to specific OA/TA neuronal subtypes. The cell-specific reagents described in this study will have a wide range of applications spanning neuroanatomy, physiology, and behavior. Functional characterization of these subtypes can provide insight into the neural logic of neuromodulation by other biogenic amines including noradrenaline in vertebrates.

### Specificity of genetic tools to manipulate identified OA/TA neurons

Genetic reagents that target specific OA/TA subtypes are crucial for dissecting the multifunctional roles of octopaminergic and tyraminergic signaling. However, access to the full repertoire of OA/TA neurons has remained incomplete. To overcome this limitation, we have established a comprehensive genetic tool to dissect the function of *Drosophila* long-range OA/TA neurons. Our work powerfully augments the toolbox to manipulate OA/TA neurons in a subtype-specific manner by two ways. First, we generated split-GAL4 lines that label 14 OA/TA subtypes that have been previously inaccessible by GAL4 drivers. Second, our split-GAL4 lines complement previously published drivers for OA/TA neurons, which were often created as part of characterizing resources for specific subregions, such as visual circuits (Nern et al., 2025), olfactory circuits (Aso et al., 2014; Shuai et al., 2025), or descending neurons (Namiki et al., 2018; Zung et al., 2025). Although these split-GAL4 lines can selectively label particular OA/TA neuronal types within each category, we found that some of these lines also exhibited expression in additional OA/TA neuronal subtypes outside the intended category. We focused on isolating GAL4 lines that label only single or at most three OA/TA types across the whole brain, thereby improving labeling specificity and increasing the number of genetic reagents available to access these cells. This level of specificity is practically important for minimizing potential transgene-specific artefacts when interrogating the functional role of individual OA/TA neurons. Intriguingly, most split-GAL4 lines selective for central VM neurons label multiple subtypes in a combinatorial manner, possibly reflecting their shared neurodevelopmental origins (Busch and Tanimoto, 2010).

### Revising OA/TA neurons classification

Detailed examinations of neurons labeled by the new split-GAL4 lines coupled with connectome data analysis have led us to propose modifications to the previously established OA/TA neuronal type classification (Busch et al., 2009). Two new cell types (ASM4, ASM5) are added as novel OA/TA neurons. Contrary, high degree of morphological and connectivity similarities between ASM2 and ASM3, as well as between VUMa6 and VUMa7, suggest these neurons form single subtypes. Neuroanatomical similarities between these two groups have been previously reported (Busch et al., 2009). Coincidentally, recent update on Codex nomenclature renamed VUMa7 as VUMa6. The fact that ASM2/3 and VUMa6/7 are often labeled together by our split-GAL4 lines provides circumstantial evidence that they share genetic characteristics. Functions of both ASM and VUMa neuronal types have been poorly characterized so far (Burke et al., 2012; Busch et al., 2009; Crocker et al., 2010; Hermanns et al., 2022; Meschi et al., 2024), but our specific split-GAL4 lines for these diverse neurons will permit new neuroanatomical, connectivity, and functional analyses.

Our data also promote re-classifications of VUMd subtypes. Our neuroanatomical and connectivity analyses suggest that VUMd1 neurons exist as a pair of asymmetrical neurons, a characteristic of VPM neurons (Busch et al., 2009). In contrast, VUMd2 neurons likely consist of two subtypes with distinct morphology and connectivity. If VUMd3 and VUMd4 neurons annotated in Codex indeed release OA/TA, each of them also likely constitutes a distinct cell type. Several recent studies have characterized subsets of OA/TA descending neurons (Babski et al., 2024; LeDue et al., 2016; Namiki et al., 2018), reflecting growing interest in these cells, which are uniquely positioned to interface with the central brain and premotor pathways. Our findings are consistent with emerging evidence suggesting heterogeneity among other VUMd neuronal subtypes (Babski et al., 2024; Szuperak et al., 2025).

Lastly, evidence supporting the classification of VPM5 and AL2b2 as OA/TA neurons remains limited, leaving open the possibility that these cells are not OA/TA neurons. VPM5 clones have been identified only a few times, and neurons with similar morphology have not been identified in any EM volumes yet (Bates et al., 2025; Berg et al., 2025; Dorkenwald et al., 2024; Schlegel et al., 2024). Additionally, the total number of all AL2 subtypes annotated in Codex (nine per hemisphere), which do not include AL2b2, slightly exceeds the number of Tdc2-expressing cells in the AL2 cluster (7-8; (Busch et al., 2009; Ishii et al., 2022)). AL2b2 is predicted to be cholinergic in the EM volumes. Characterization of these cell types requires further experimentation.

### Specific ASM subtypes suppress social isolation-dependent aggression

Although cell-type specific functions of OA/TA subtypes have been suggested for more than a decade (Burke et al., 2012; Certel et al., 2010; Crocker et al., 2010; Zhou et al., 2008), experimental validation has been limited to a handful of cases (Hermanns et al., 2022; Kapoor and Waddell, 2024; LeDue et al., 2016; Sayin et al., 2019). In these studies, the number of specific cell types under investigation were limited by the availability of genetic reagents. As a result, the contribution of other OA/TA subtypes could not be examined, leaving open the possibility that multiple OA/TA neurons cooperatively modulate the physiological processes or behaviors under study. A comprehensive toolkit for manipulating all OA/TA subtypes in parallel will greatly facilitate efforts to determine whether specific physiological or behavioral functions of OA/TA arise from individual neuronal subtypes or from coordinated activity across groups of OA/TA neurons.

Indeed, by using the newly generated split-GAL4 lines, we were able to determine specific OA/TA neuronal subtypes that influence two important behaviors: visual flight control (discussed below) and aggressive behavior. Using these tools, we identified two specific subtypes of ASM neurons (ASM4 and ASM5) that suppress aggression in both males and female flies. Experience-dependent modulation of aggressive behavior is crucial to prevent maladaptive aggression (Hsu et al., 2006). However, despite its clear ethological importance, the neuronal mechanisms that regulate this process remain poorly understood. Our previous work showed that a *nvy*-expressing OA/TA subset includes neurons that suppress social-dependent aggression in both sexes (Ishii et al., 2022). The majority of *nvy*-expressing OA/TA neurons fall within the ASM and VL clusters (Ishii et al., 2022). These new split-GAL4 lines enabled us to identify ASM4/5, among the seven subtypes encompassed within these two clusters, as the neurons responsible for suppressing aggression among socially isolated flies. Although current tools prevent us from clearly separating ASM4 and ASM5, the aggression-suppressing phenotype produced by the activation of OTA-SG10, which labels ASM4 more consistently than ASM5, implies that ASM4 may be more relevant for modulating aggression. Like most other OA/TA neurons, ASM4/5 neurons are sexually monomorphic (Fig. 1E, F; (Berg et al., 2025)). The neural mechanism through which these neurons modulate sexually dimorphic aggressive behavior is an important topic for future study.

### Layer-specific innervation by AL2 neurons implement precision modulation of visual circuits

Anatomical characterization of AL2 neurons reveals a striking layer specificity of neuromodulatory innervation in the optic lobe (Fig. 2G, H). Distinct visual features such as color, luminance change, motion, optic flow and feature detection are processed *via* anatomically segregated pathways (Ryu et al., 2022). Layer specificity results from selective synaptic connectivity. The FlyWire annotation of FAFB shows chemical synaptic connections between AL2 and neurons that have been shown to play a key role in optomotor stabilization and object tracking behaviors. AL2i1 forms synaptic connections with T4 and T5 neurons, the primary directionally selective motion detecting neurons in the visual system that are thought to drive optomotor gaze stabilization (Busch et al., 2018; Dorkenwald et al., 2022; Haikala et al., 2013; Joesch et al., 2010; Schlegel et al., 2024; Strother et al., 2017). By contrast, AL2i2 forms synaptic connections with T3 neurons, which are omnidirectional feature detectors shown to be important for object tracking behavior (Dorkenwald et al., 2024; Frighetto and Frye, 2023; Schlegel et al., 2024). Accordingly, optogenetic activation of AL2i2 (OTA-SG11), and presumably gain modulation of T3, enhanced bar tracking behaviors, but did not significantly alter optomotor performance (Fig. 7C_2_, D_2_, E_2_). By contrast, activation of AL2i1 (SS32637), and presumably neuromodulation of T4/T5, saturated optomotor performance toward the maximum PI=1; (Fig. 7C_1_, D_1_) and significantly increased saccade rate during optomotor stabilization behavior (Fig.7E_1_), without changing the overall performance of bar tracking (Fig. 7C_2_, D_2_). By contrast, constitutive hyperpolarization of AL2i1 results in reduced optomotor performance (Fig. 7C_3_, D_3_). These results are consistent with work showing that calcium accumulation by Tdc2-positive OA/TA neurons is coupled to the transition from quiescence to the onset of flight, and that this response is required for the locomotion-induced increase in response gain by directionally selective visual neurons (Strother et al., 2018; Suver et al., 2012).

Our findings suggest that the locomotion dependent increase in response gain of directionally selective neurons, and visual stabilization reflexes, is mediated by OA/TA neuromodulation of T4/T5 neural pathways by AL2i1 neurons in particular, whereas flexible modulation during flight of object tracking can be achieved by modulation of T3 by AL2i2 neurons. With novel OA/TA split-Gal4 lines, these hypotheses are now directly testable with *in-vivo* calcium imaging coupled with optogenetics. The selective action of AL2 classes is remarkable since it is generally thought that neuromodulation acts broadly through volume release (Marder, 2012; Nässel, 2009; van den Pol, 2012). To our knowledge, ours is the first demonstration that OA/TA neuromodulatory neurons can selectively influence distinct local visual circuits and their respective behaviors.

### The application of the OA/TA genetic tools

In addition to cell type-specific manipulations, genetic reagents described in this study can provide crucial support to the collective effort of rigorously annotating the fly connectome. Connectomes are transforming neuroscience and, to date, *Drosophila melanogaster* has one of the most complete single-neuron-based circuit maps of the brain (Bates et al., 2025; Berg et al., 2025; Dorkenwald et al., 2024; Scheffer et al., 2020; Schlegel et al., 2024; Zheng et al., 2018). Confocal microscopy images provide information to generate high-resolution maps of neuronal connections by complementing neuronal volumes reconstructed from EM images. Connectome analysis can also provide testable hypotheses about how OA/TA neurons might exert effects on behavior and physiology. For example, whereas we did not observe gross sexual dimorphism among OA/TA neurons labelled by available lines, connectomic analysis reveals sex differences in connectivity and synaptic distributions in addition to morphology (Berg et al., 2025).

Several limitations of the current set of split-GAL4 lines must be acknowledged. First, several known classes of OA/TA neurons remain inaccessible by the split-GAL4 approach despite our attempts. They include VUMd3 and VUMd4 neurons, as well as the AL1 cell type (Busch et al., 2009; Busch and Tanimoto, 2010). In addition, we did not attempt to isolate split-GAL4 lines for OA/TA neurons that have cell bodies in the VNC. OA/TA neurons are important for *Drosophila* motor functions, including locomotion, flight, and copulation (Babski et al., 2024; Brembs et al., 2007; Rezával et al., 2014; Schützler et al., 2019). OA/TA descending neurons can contribute to these functions, but it is likely that OA/TA neurons intrinsic to the VNC have direct impact on the physiology of premotor and motor neurons, thus affecting motor output of ethologically important behaviors. Another important limitation is that we have not determined whether the different subtypes release OA or TA (or both). Previous studies show that subsets of Tdc2-expressing neurons, including VL and ASM subtypes, are not always immunoreactive with Tbh or octopamine itself (Burke et al., 2012; Busch et al., 2009; Zhou et al., 2008).This is an important topic for future studies, as OA and TA are known to have distinct and sometimes antagonistic effects in the insect nervous system (Brembs et al., 2007; Huang et al., 2016; Ishikawa et al., 2016; Roeder, 2005; Schützler et al., 2019). Lastly, it is necessary to use multiple split-GAL4 lines to confirm subtype-specific effects because most split-GAL4 lines label multiple cell types, within and outside the OA/TA neurons. Although the specificity of split-GAL4 lines is generally superior to conventional GAL4 lines, they usually do not limit the expression to a single class of neurons.

Together, our results establish one of the most comprehensive toolsets for the cellular and genetic dissection of a specific biogenic amine neuronal class in any model organism to date. By integrating precise genetic access with anatomical and behavioral analyses, we establish a framework to systematically map aminergic neuromodulatory circuits. Beyond their relevance for invertebrates, the *Drosophila* OA/TA system offers a powerful comparative model for understanding how small populations of neuromodulatory neurons achieve multifunctionality - a principle that may extend to noradrenergic systems across the animal kingdom.

## Supporting information

Raw Data File

## Acknowledgements

This work has been supported by NIH NIMGS R35GM119844 to K.A., and NIH NINDS R01NS120984 to M.A.F.

## Author contributions

L.M.P.C. conceptualization, investigation, validation, formal analysis, visualization, writing; V.F. data curation, formal analysis, investigation, methodology, validation, visualization, writing; E.Y. investigation, validation, visualization; B.R. formal analysis, visualization; S.S. investigation, validation; V.L. investigation; A.K. investigation; J.H. investigation, validation; K.I. investigation; V.M. investigation; M.T. investigation; C.T.W: investigation; G.F. formal analysis; M.A.F.: conceptualization, funding acquisition, project administration, resources, supervision, writing; K.A. conceptualization, data curation, formal analysis, funding acquisition, project administration, resources, supervision, visualization, writing.

## Materials and Methods

### Key Resource Table

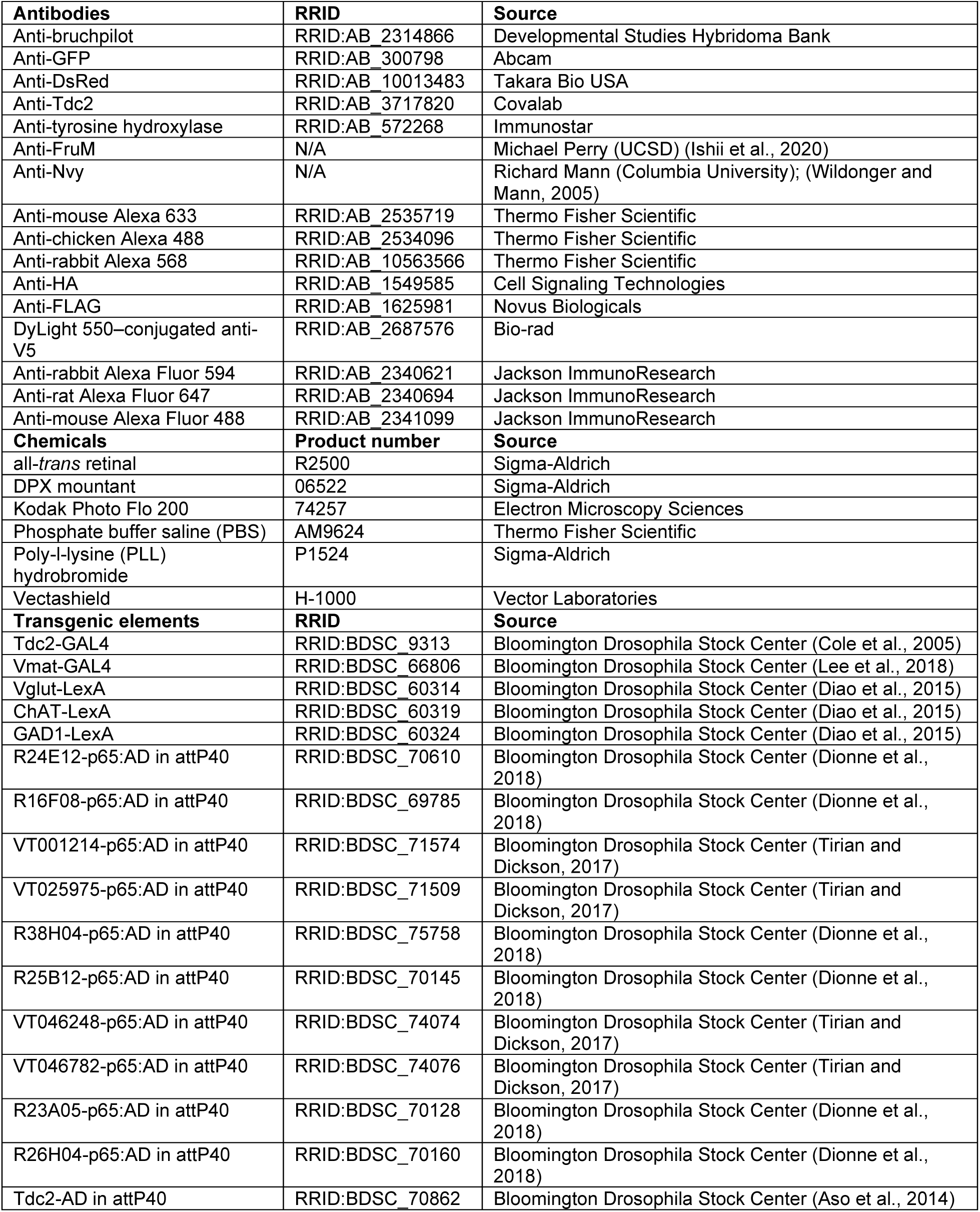
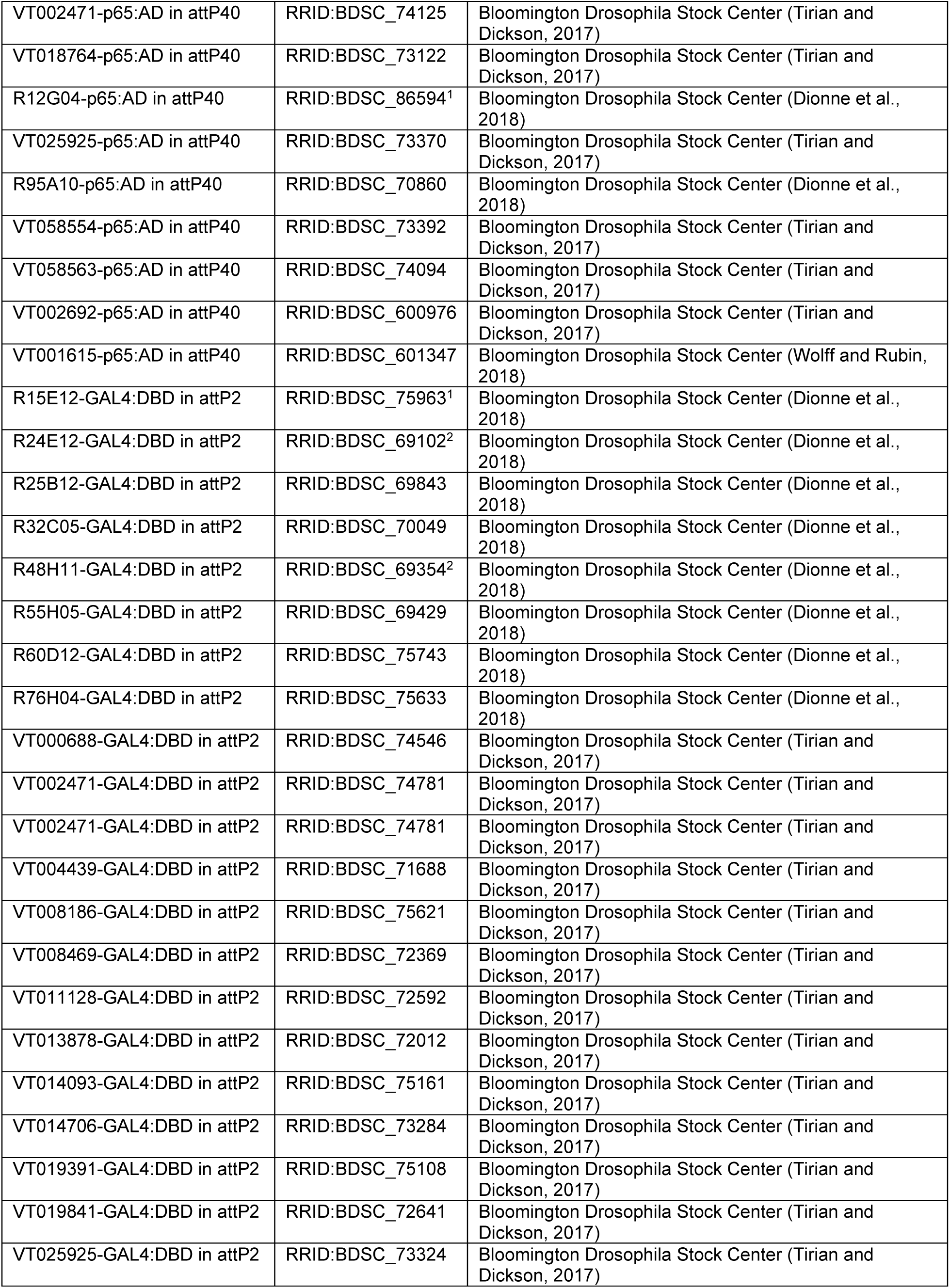
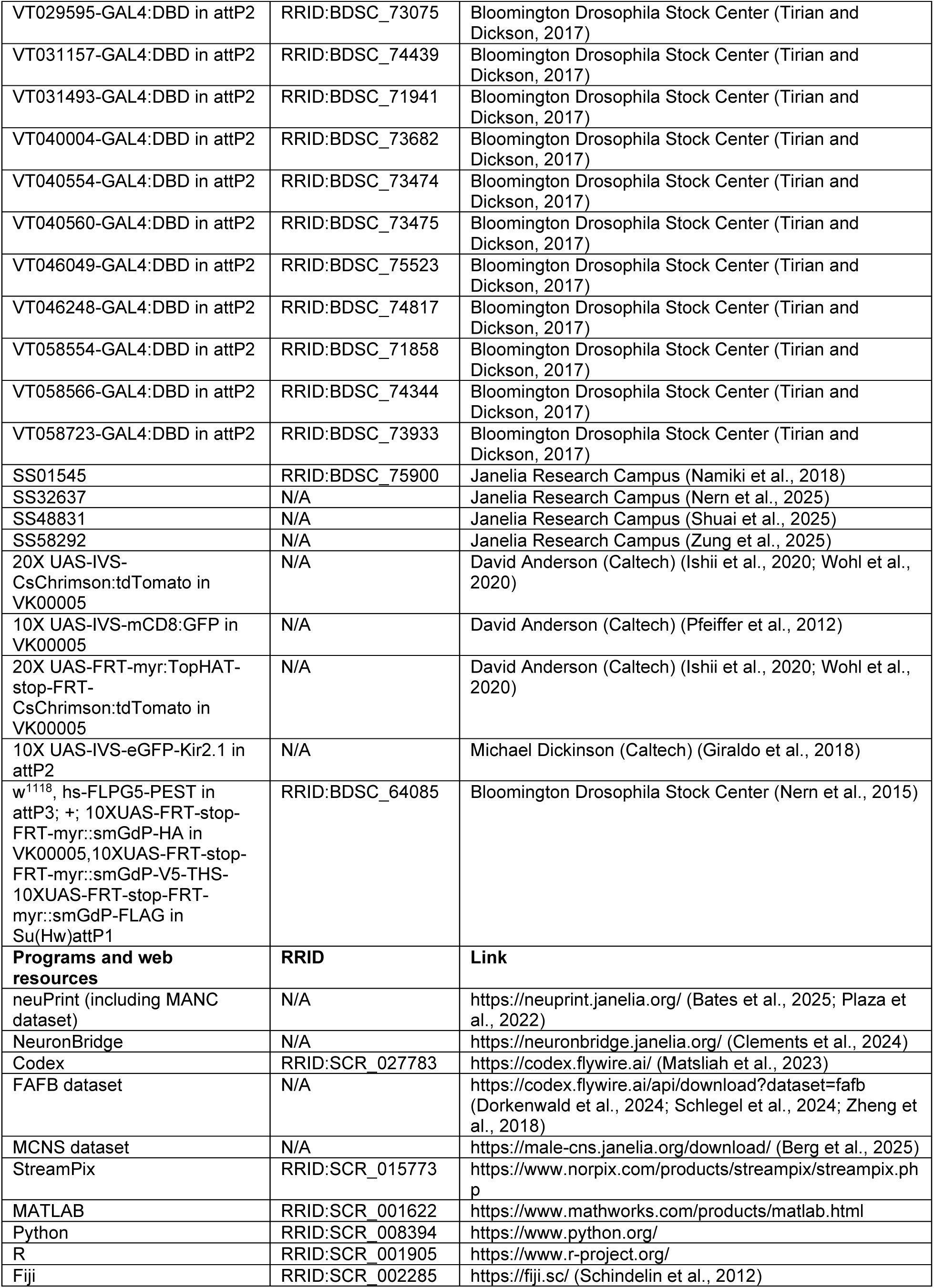
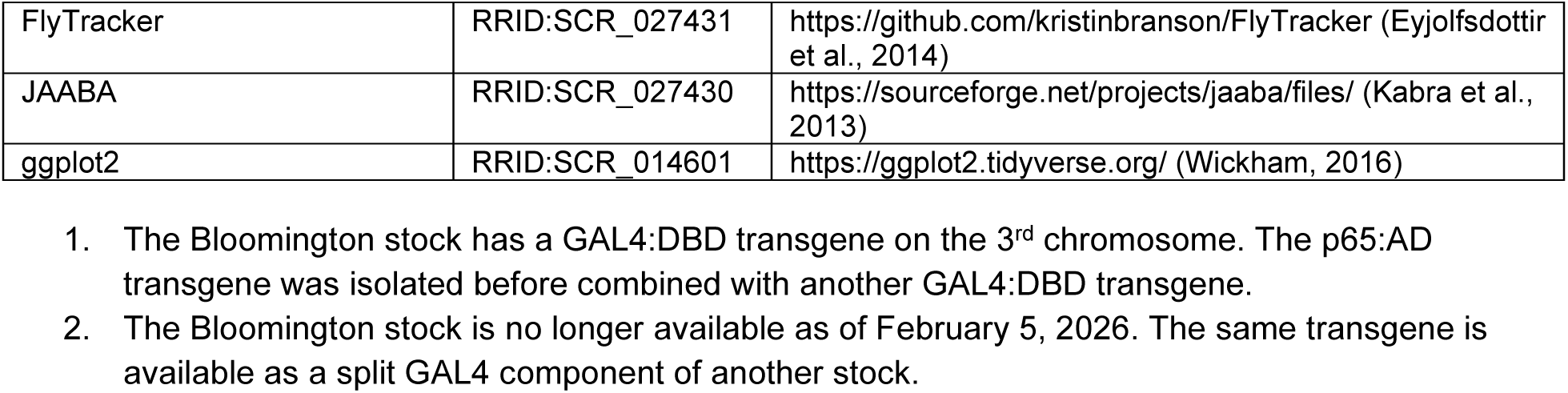

### Drosophila strains

The sources of the *Drosophila* strains used in this study are indicated in the Key Resource Table. The complete list of transgenic strains is provided in Supplementary Table S1.

### Selection of hemidriver lines

Several methods were used to identify the transgenic lines that express either the activation domain (AD) or DNA-binding domain (DBD) of the split-GAL4 protein. 1) We first manually scanned the images of the first generation GAL4 lines of the FlyLight collection to find candidate promoter fragments that might label OA/TA neurons. Hemidrivers were combined with either *Tdc2-AD* (Aso et al., 2014) or with one another. 2) Next, we screened for the GAL4 lines that likely contain the registered neural traces of known OA/TA neurons in the hemibrain EM volume. The EM traces in Neuprint platform were used as search queues in NeuronBridge for GAL4 lines with overlapping labeling patterns with the queue volumes. 3) Lastly, we used the single cell clone images from *Tdc2-GAL4* line as the queue to directly search for candidate GAL4 lines using NeuronBridge. The OA/TA neuronal clone was first generated by MCFO (Nern et al., 2015) as described below. The images with successful registration to the standard *Drosophila* brain volume were used to identify GAL4 lines with overlapping expression pattern.

### Screening process of split-GAL4 lines

We combined candidate AD and DBD hemidrivers to create split-GAL4 lines. Males of these lines were crossed with females carrying *20X UAS-CsChrimson:tdTomato* (in VK00005) (Ishii et al., 2020) to visualize the neuronal processes. CsChrimson was preferred because of its robust localization to fine neuronal processes. Expression patterns in the brain of 2-3 male offspring per cross was first analyzed. Lines that appeared to label putative OA/TA neurons were examined again with increased number of brains for consistency and identification of labeled cell types. Lines with 1) consistent labeling pattern, and 2) relatively sparse labeling were further selected for analysis of expression patterns in brains of both sexes and in male VNC, as well as co-labeling with anti-Tdc2 antibody.

### Immunohistochemistry

#### Conventional immunostaining

Fly brains were dissected in cold phosphate buffer saline (PBS, Thermo Fisher Scientific, cat# AM9624) and fixed in 4% formaldehyde in 1X PBS at room temperature for 15 minutes in 6 × 10 microwell minitray (Thermo Fisher Scientific, cat# 439225). Brains were washed in PBST (0.3% Triton X-100 in 1X PBS) for 10 min X 3 times in a rotating 1.5mL tubes, followed by incubation in a blocking solution (10% heat-inactivated normal goat serum in 1X PBS) for 30 min, without rotating. Primary antibodies diluted with the blocking solution (1:10 (supernatant) or 1:100 (concentrated) for mouse anti-bruchpilot (BRP) (Developmental Studies Hybridoma Bank nc82, RRID: AB_2314866), 1:200 for mouse anti-tyrosine hydroxylase (Immunostar, cat# 22941, RRID: AB_572268), 1:1000 for chicken anti-GFP (Abcam, cat# ab13970, RRID: AB_300798), 1:1000 for rabbit anti-DsRed (Takara Bio USA, cat# 632496, RRID: AB_10013483), 1:100 for guinea pig anti-Fru^M^ (gift from Michael Perry, University of California, San Diego) (Ishii et al., 2020), 1:200 for rabbit anti-Tdc2 (Covalab, cat# 00013520, RRID: AB_3717820) and 1:1000 for rabbit anti-Nvy (gift from Richard Mann, Columbia University) (Wildonger and Mann, 2005) were applied to the samples in 1.5mL tubes at 4°C for 2 days. The brains were washed in PBST for 10 min X 3 times in rotating 1.5mL tubes, followed by incubation in a blocking solution for 30 min, without rotating. Brains were then transferred in 6 × 10 microwell minitray and incubated in secondary antibodies diluted with the blocking solution (1:100 for goat anti-mouse Alexa 633 (Thermo Fisher Scientific, cat# A-21052, RRID: AB_2535719), 1:100 for goat anti-chicken Alexa 488 (Thermo Fisher Scientific, cat# A-11039, RRID: AB_2534096), and 1:100 for goat anti-rabbit Alexa 568 (Thermo Fisher Scientific, cat# A-11036, RRID: AB_10563566)) at 4°C overnight. Brains were washed in PBST for 10 min X 3 times and then incubated in the 50% glycerol/PBS at room temperature for 2 hours, followed by 75% glycerol/PBS at room temperature for 1.5 hours. Samples were mounted in Vectashield (Vector Laboratories, cat# H-1000) onto a glass slide.

#### Immunostaining for characterizing neuroanatomy of split-GAL4 lines

For OA/TA split-GAL4 lines characterization, fixation, dehydration, and mounting processes of the “DPX mounting” protocol developed by HHMI Janelia Research Campus (www.janelia.org/project-team/flylight/protocols) were integrated with a conventional immunostaining protocol. Unless otherwise specified, all steps were carried out in 6 × 10 microwell minitray. Fly brains were dissected in cold Schneider’s Insect Medium (S2) Schneider’s medium and fixed in 2% formaldehyde in Schneider’s medium at room temperature for 1 hour. Brains were washed in PBST for 5 min X 4 times, followed by incubation in a blocking solution for 30 min. Primary antibodies diluted with the blocking solution (1:10 (supernatant) or 1:100 (concentrated) for mouse anti-bruchpilot (BRP) (Developmental Studies Hybridoma Bank nc82, RRID: AB_2314866), 1:1000 for rabbit anti-DsRed (Takara Bio USA, cat# 632496, RRID: AB_10013483) were applied to the samples at 4°C for 1-2 days. The brains were washed in PBST for 10 min X 3 times and then incubated in secondary antibodies diluted with the blocking solution (1:100 for goat anti-mouse Alexa 633 (Thermo Fisher Scientifi, cat# A-21052, RRID: AB_2535719), 1:100 for goat anti-chicken Alexa 488 (Thermo Fisher Scientific, cat# A-11039, RRID: AB_2534096), and 1:100 for goat anti-rabbit Alexa 568 (Thermo Fisher Scientific, cat# A-11036, RRID: AB_10563566)) at 4°C overnight.

After incubation with secondary antibody, brains were washed in 0.5% PBT for 30 min X 3 times and then incubated in post-fixation solution (4% formaldehyde in PBS1X) for 4 hours at RT, followed by 0.5% PBT washes for 15 min X 4 times.

Coverslips (22 mm by 22 mm square no.1, Globe Scientific Inc., cat# 1404-10) were dipped twice in a coating solution (0.08% (w/v) poly-l-lysine (PLL) hydro- bromide (Sigma-Aldrich, cat# P1524) and 0.2% Kodak Photo Flo 200 (Electron Microscopy Sciences, cat# 74257) in water, stored at 4°C) and air-dried one day before mounting. Slides with spacers were made as described in the above website. Before mounting, brains were washed in 1X PBS for 5 min X 2 times to remove detergents. Brains were transferred on the PLL-coated coverslip and gently placed on the glass surface with the anterior part facing down. After transferring all brains, the coverslips were briefly washed with water to remove PBS and then subjected to dehydration series by 10-min soaks in successive baths of ethanol solutions (30, 50, 75, 95, 100, 100, and 100%), followed by 5-min soaks in three successive baths of xylene for clearing. The coverslips taken out of xylene were held horizontally and applied immediately with 200 μl of DPX mountant (Electron Microscopy Sciences, cat# 13512). The coverslips were flipped over the slides and placed between the spacers and gently pressed down once. The DPX-mounted slides were air-dried in the dark for 2 to 3 days before microscopic observations.

#### Single-cell stochastic labeling

Single-cell stochastic labeling was performed following the Multi Color Flip Out (MCFO) approach described in (Nern et al., 2015). Virgin females of the MCFO maternal strain (RRID:BDSC_64085) were crossed with Split-GAL4 males of the following lines: OTA-SG08, OTA-SG09, OTA-SG17, SG18, OTA-SG23, OTA-SG25, OTA-SG27, OTA-SG30, OTA-SG32, OTA-SG37, OTA-SG38, OTA-SG39. The F1 male offspring with the following genotype were used for the MCFO experiment: *w, hs-FLPG5-PEST in attP3/Y; AD-GAL4/+; 10XUAS-FRT-stop-FRT-myr::smGdP-HA in VK00005,10XUAS-FRT-stop-FRT-myr::smGdP-V5-THS-10XUAS-FRT-stop-FRT-myr::smGdP-FLAG in Su(Hw)attP1/ DBD-GAL4*.

F1 male offspring used for MCFO experiment were reared at 25°C from embryo to adult. 6-7 days after eclosion flies were warmed at 37°C in a water bath (heat shock) for 10 to 20 min, and then kept at 25°C for at least 4 days before dissection. For each Split-GAL4 driver was performed preliminary investigation of the optimal heat shock conditions to obtain sparse labelling. Optimal heat-shock time used are: 10 min for OTA-SG37 and SG39, 12 min hs for OTA-SG08 and SG09, 15 min for OTA-SG17, 20 min for OTA-SG18 and SG30. Initial heath-shock testing (from 12 to 40 min) did not yield any clones in the following lines: OTA-SG23, SG25, SG27 and SG32.

Subsequent immunostaining steps were performed according to the protocol described in “Immunostaining for characterizing neuroanatomy of split-GAL4 lines” above. Briefly, brains were dissected in ice-cold Schneider’s medium and fixed in 2% formaldehyde in Schneider’s medium at room temperature for 60 min, followed by 0.5% PBT washes for 10 min X 4 times. Samples were incubated in a blocking solution (5% goat serum/0.5%PBT) at room temperature for 90 min. Primary antibodies (1:300 dilution of anti-HA rabbit monoclonal antibody (mAb) (C29F4, Cell Signaling Technologies, cat# 3724S, RRID:AB_1549585), 1:200 dilution of anti-FLAG rat mAb (DYKDDDDK epitope tag antibody (L5), Novus Biologicals, cat# NBP1-06712, RRID:AB_1625981), and 1:10 dilution of supernatant anti-bruchpilot mouse mAb (Developmental Studies Hybridoma Bank nc82, RRID:AB_2314866), all diluted in 5% goat serum/0.5%PBT) were applied at 4°C for 2 days. The brains were washed in 0.5%PBT for 30 min X 3 times and then incubated with secondary antibodies (1:500 dilution of anti-rabbit Alexa Fluor 594 (Jackson ImmunoResearch, cat# 711-585-152, RRID:AB_2340621), 1:150 dilution of anti-rat Alexa Fluor 647 (Jackson ImmunoResearch, cat# 712-605-153, RRID:AB_2340694), and 1:150 dilution of anti-mouse Alexa Fluor 488 (Jackson ImmunoResearch, cat# 715-545-151, RRID: AB_2341099), all diluted in 5% goat serum/0.5%PBT) for two days at 4°C. Brains were washed in 0.5%PBT for 30 min × 3 and then incubated with DyLight 550–conjugated mouse anti-V5 (Bio-Rad, cat# MCA1360D550GA, RRID: AB_2687576) (1:500 dilution in 5% goat serum/0.5%PBT) at 4°C overnight. Processes for washout of the secondary antibodies, brain clearing, and mounting were identical to the protocol above.

#### Fischbach sample preparation

Brains were dissected and stained as described above. Following secondary antibody incubation and washout, brains were transferred to mounting medium (Vectashield, Vector Laboratories, cat# H-1000) for 10 min at room temperature. For mounting, each brain was positioned dorsally, with the ventral surface in contact with the microscope slide. Coverslips were prepared in advance by placing a small amount of clay at each corner (approximately the size of a *Drosophila* brain) to serve as spacers. The prepared coverslip was gently lowered onto the sample, and once the correct dorsal orientation was confirmed, it was lightly pressed at the corners to ensure that the ventral surface contacted the slide while the dorsal surface was secured beneath the coverslip.

#### Image acquisition and post-processing

Images were acquired by a Zeiss LSM 700 (UCLA), LSM 880 (at the Waitt Advanced Biophotonics Core, Salk Institute) or Zeiss LSM900 (shared in Molecular Neuroscience Department at Salk Institute). Whole-brain z-stack images were acquired using a 20X air objective, while high-resolution images from Fischbach samples were acquired using a 40X oil-immersion objective. Maximum z-projections were generated on Fiji software (RRID: SCR_002285; https://fiji.sc/). Fiji was also used to crop the original images to standardize the sample appearance, and to adjust signal intensities to enhance the visibility.

### Connectivity similarity analysis

The raw synapse count data of FAFB, MCNS, and MANC was downloaded from the following links:

FAFB: https://codex.flywire.ai/api/download?dataset=fafb
MCNS: https://male-cns.janelia.org/download/#__tabbed_1_2
MANC (v1.2.1): https://connectome-neuprint.github.io/neuprint-python/docs/index.html

Cell type definition is based on FAFB v783. Only synapses that belong to traced and annotated neurons were considered. Upstream and downstream neurons in MCNS and MANC that have less than 5 synapses with the given OA/TA neuron were excluded from the analysis, following the criteria of Codex synaptic partners. For each OA neuron, the synapse counts from upstream cell types and to downstream cell types were concatenated into one vector representing the upstream and downstream connectivity information. The pairwise cosine similarity of connectivity between two OA/TA neurons (S) is defined as the dot product of the connectivity vectors of two neurons of interest (A and B) divided by the product of A and B’s magnitudes, as calculated by the following equation:

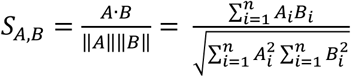

Where n is the number of total downstream or upstream neurons for all OA/TA neurons.

Due to relatively limited synapse annotations, we did not use BANC (brain and nerve cord) and FANC (female nerve cord) EM volumes for the connectivity analysis.

### Behavioral assays

#### Assays for aggressive behavior

Male and female flies were collected on the day of eclosion and isolated for 6 days before the assay. From day 4 flies were reared on food supplied with 0.2 mM all-*trans* retinal (Sigma-Aldrich, cat# R2500, 20 mM stock solution prepared in 95% ethanol) supplemented with 5% yeast, and the vials were covered with aluminum foil to avoid light exposure. Behavior experiments were conducted in the late afternoon (ZT10-ZT13) at 25°C with 60% relative humidity in climate-controlled booths, largely described as in (Wohl et al., 2024). Briefly, the constant infrared (850nm) illumination and timed red light (655nm) for optogenetic stimulation were provided by custom-designed multicolor LED backlights (Metaphase Technologies Inc). Videos were acquired using Streampix software (version 8.5.0.0, Norpix) in MP4 format using H.264 GPU-accelerated real-time compression. Flies were discarded after each experiment, and the behavior devices were remade with fresh food substrate after every recording.

To optogenetically activate neurons that transgenically express CsChrimson, 655nm red-light was delivered in 10ms pulses at 5Hz at an intensity of 12μM/cm^2^ using the above-mentioned multicolor LED backlights. Optogenetic stimulation periods were custom-scripted and controlled from Streampix for acquisition synchronized stimulation: ON/OFF TTL output from StreamPix were sent to an Auduino Uno R3 microcontroller board via a USB-6001 DAC (part #782605-01, National Instruments), which then outputs 10msec ON/OFF TTL signals at the specified rate (i.e. 5Hz) to the LED controller (Metaphase Tech ULC-2).

Behavior movies were analyzed as described in (Leng et al., 2020). Briefly, the movies were first processed by FlyTracker (Eyjolfsdottir et al., 2014); (https://github.com/kristinbranson/FlyTracker) and JAABA (Kabra et al., 2013) (https://sourceforge.net/projects/jaaba/files/). A JAABA-based lunge classifier and a custom script of smoothening filter were used to detect lunges for male flies and headbutt for female flies (Leng et al., 2020).

#### Assay for visual flight behavior

##### Animal preparation

Progeny from crosses between split-GAL4 driver lines and UAS-CsChrimson were transferred to media containing 0.5mM of all-*trans* retinal (Sigma-Aldricb, cat# R2500) upon eclosion,and were reared in darkness for 3-5 days before experimentation to prevent premature activation. Flies were prepared for visual flight assays as previously described in (Frighetto and Frye, 2023). Briefly, they were cold-anesthetized on a Peltier cell at ∼4°C. Stainless-steel pins (Fine Science Tools, cat# 26002-10; ∼1 cm) were attached to the dorsal thorax using UV-activated glue (Esslinger, cat# 12.201), producing a body angle of ∼30°. Flies recovered upside-down for at least 30 min before experimentation and were provided with small pieces of paper to cling to, preventing flight and conserving energy.

##### Magnetic-tether setup

Visual stimuli were presented using a cylindrical LED display spanning 360° in azimuth and 56° in elevation (460/243 nm, spectral peak/FWHM) as described previously (Duistermars and Frye, 2008; Reiser and Dickinson, 2008). Each LED subtended ∼3.75° on the fly’s retina at the horizontal midline. Flies were suspended between two magnets, allowing free rotation about the yaw axis. In this virtual reality visual flight simulator, the freely revolving animal has true closed-loop feedback control over its own visual and proprioceptive experience. Six infrared diodes (940 nm peak emission) illuminated the fly from above, and behavior was recorded from below at 100 frames s⁻¹ using an infrared-sensitive camera (Blackfly S USB3, Teledyne FLIR, cat# BFS-U3-04S2C-CS BFS-U) fitted with a zoom lens (InfiniStix 1.0×/94 mm, Infinity Photo-Optical, Edmund Optics cat# 55-359) and a long-pass filter (Thorlabs, cat# FGL850M). Each trial began with 10s of acclimation followed by 20s of panoramic motion (clockwise and counterclockwise, 120° s⁻¹) to elicit an optomotor response. Flies that failed to complete this trial or showed excessive wobble were excluded.

##### Optogenetics

Optogenetic activation was achieved using a fiber-coupled red LED (Thorlabs, cat# M617F2; 617nm) combined with a long-pass dichroic mirror (Thorlabs, cat# DMLP805R and CM1-DCH) to reflect red light for stimulation and transmit infrared light for imaging. An aspheric lens (Thorlabs, cat# AL50100M) collimated the red beam, which was directed from below the tethered fly and focused to cover the whole fly (∼300µm in diameter).

##### Visual stimulation

Visual stimuli consisted of either a wide-field panorama (360° × 56°) composed of randomized bright and dark vertical stripes (for gaze stabilization assay) or a motion-defined bar (30° × 56°) presented on a randomized panorama (for the object orientation assay). These stimuli rotated around the fly either singly (clockwise [CW] or counterclockwise [CCW] at 112.5° s⁻¹) or simultaneously, such that one rotated in one direction (e.g., CW bar) while the other rotated in the opposite direction (e.g., CCW panorama). Motion-defined bars were used to evoke tracking responses while minimizing luminance cues that could elicit static positional behavior. Stimulus type and speed were selected based on previous characterizations (Frighetto and Frye, 2023; Mongeau et al., 2019; Mongeau and Frye, 2017) Stimuli were generated using custom MATLAB scripts (to be made available on OSF upon publication).

Each trial lasted 10s and was followed by a 10s rest period in darkness. Each fly experienced six randomized trial conditions (3 stimuli × 2 directions). For optogenetic experiments, the six trials were first presented without stimulation, then repeated with continuous optogenetic activation throughout each trial, and finally repeated again with the stimulation off, yielding a total duration of ∼8 min. For silencing experiments, the stimulation phase was omitted, and the six trials were simply repeated twice. Only flies that flew continuously or paused briefly once during the experiment were included in the analysis.

##### Data analysis

The performance index (PI) is defined as the ratio of the fly’s angular displacement of the visual stimulus at the end of one trial. A value of 1 indicates steering that perfectly tracks the stimulus, whereas a value of −1 reflects tracking in exactly the opposite direction from the stimulus, and PI = 0 would indicate no visual response. Occasionally flies overshoot the stimulus, and PI exceeds 1. Data from magnetic-tether experiments were analyzed in R Studio using custom R scripts. Video recordings were imported into MATLAB, and fly headings were extracted using custom software. Body saccades were detected as described previously by (Mongeau and Frye, 2017). Data visualization was performed in R using the package ggplot2 (Wickham, 2016).

### Statistical Analysis

Wilcoxon signed rank test (‘signrank’ in MATLAB) was used to compare the difference of lunge numbers between different LED stimulation windows (Fig. 6B, C) within a genotype, while Mann-Whitney U-test (‘ranksum’ in MATLAB) was used to compare lunge numbers between genotypes. For the statistical analyses of visually guided behavior (Fig. 7C-E), Student’s t-test was used. Bonferroni correction of p-values for multiple comparison was applied where appropriate.

## Code Availability

Custom Python codes used for connectivity similarity analyses (Fig. 5, Supplementary Fig. S5) are available at https://github.com/asahinak/connectivity_similarity_maps. Custom MATLAB codes and a JAABA-based lunge classifier (Leng et al., 2020) are available at https://doi.org/10.6075/J0QF8RDZ.

---

**Supplementary Figure S1.**
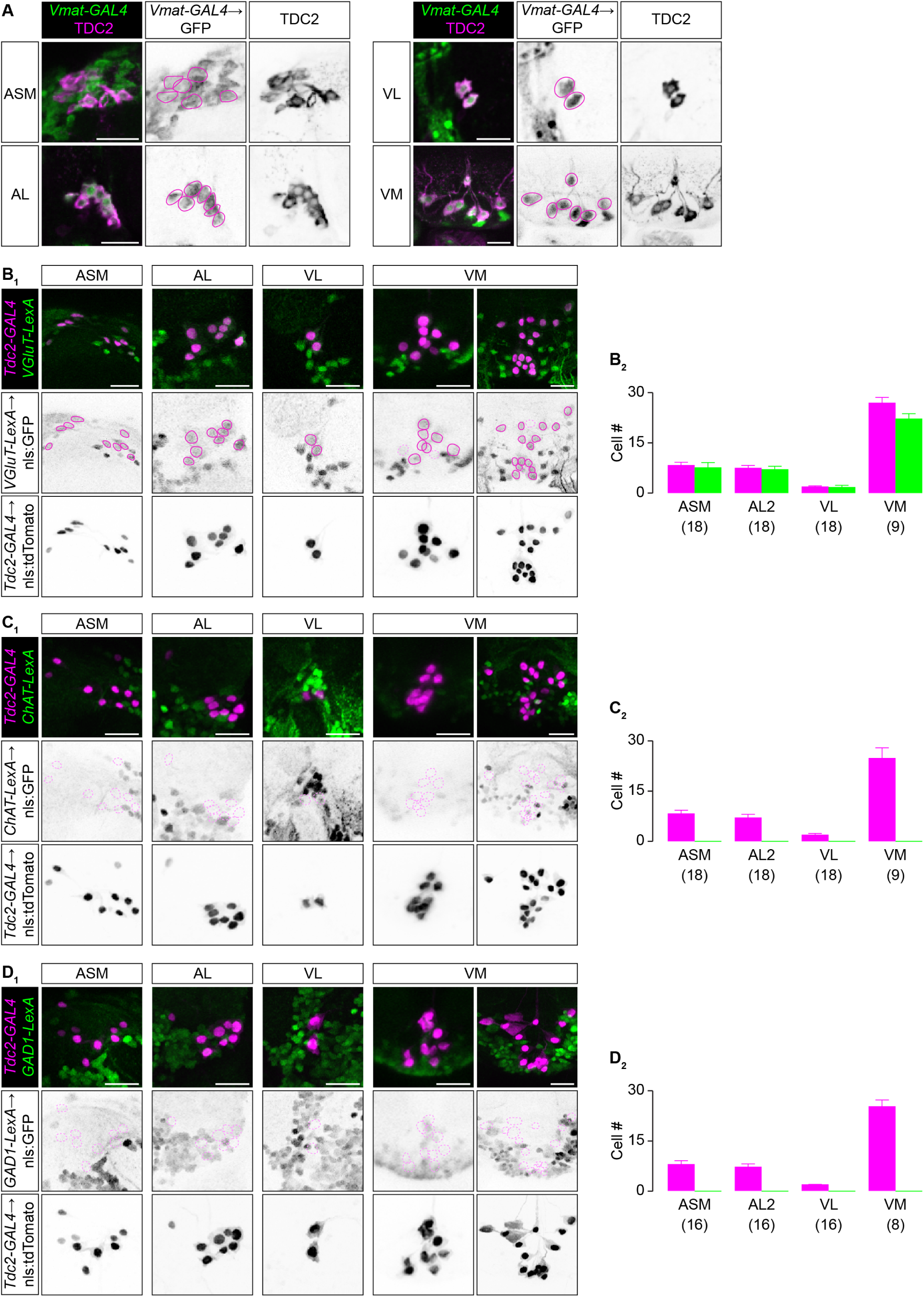
related to Figure2 1-4: OA/TA neurons co-release glutamate. (**A**) Representative OA/TA neurons from each cluster visualized by TDC2 immunoreactivity (left panels, magenta; right panels, black). GFP expressed under the control of *Vesicular monoamine transporter* promoter (*Vmat-GAL4*, left panels, magenta; middle panels, black) co-localizes with TDC2-positive neurons. (**B-D**) Immunohistochemical analysis of the overlap between OA/TA neurons, which is visualized by nuclear-localizing tdTomato expressed under Tdc2-GAL4, and GFP expressed under the LexA “knock-in” alleles of *Vescicular glutamate transporter* (*VGluT*) gene (**B**), Choline acetyltransferase (*ChAT*) gene (**C**), and Glutamic acid decarboxylase 1 (*GAD1*) gene (**D**), respectively. tdTomato was visualized by the anti-dsRed antibody (**B_1_-D_1_**, top panels, magenta; bottom panels, black), while GFP was visualized by the anti-GFP antibody (**B_1_-D_1_**, top panels, green; middle panels, black). Scale bar = 20 μm. Locations of tdTomato-expressing somata are indicated in mageta lines in the middle panel. The total number of tdTomato-expressing cells in each of four OA/TA clusters are shown in **B_2_-D_2_** (magenta), while number of cells in each cluster that co-express GFP under *VGluT-LexA* (**B_2_**), *ChAT-LexA* (**C_2_**), and *GAD1-LexA* (**D_2_**), are shown in green. Bar = mean, whisker = S.D. The number of hemispheres (for ASM, AL2, VL) or the whole brain (VM) analyzed are shown at the bottom of each bar.

**Supplementary Figure S2.**
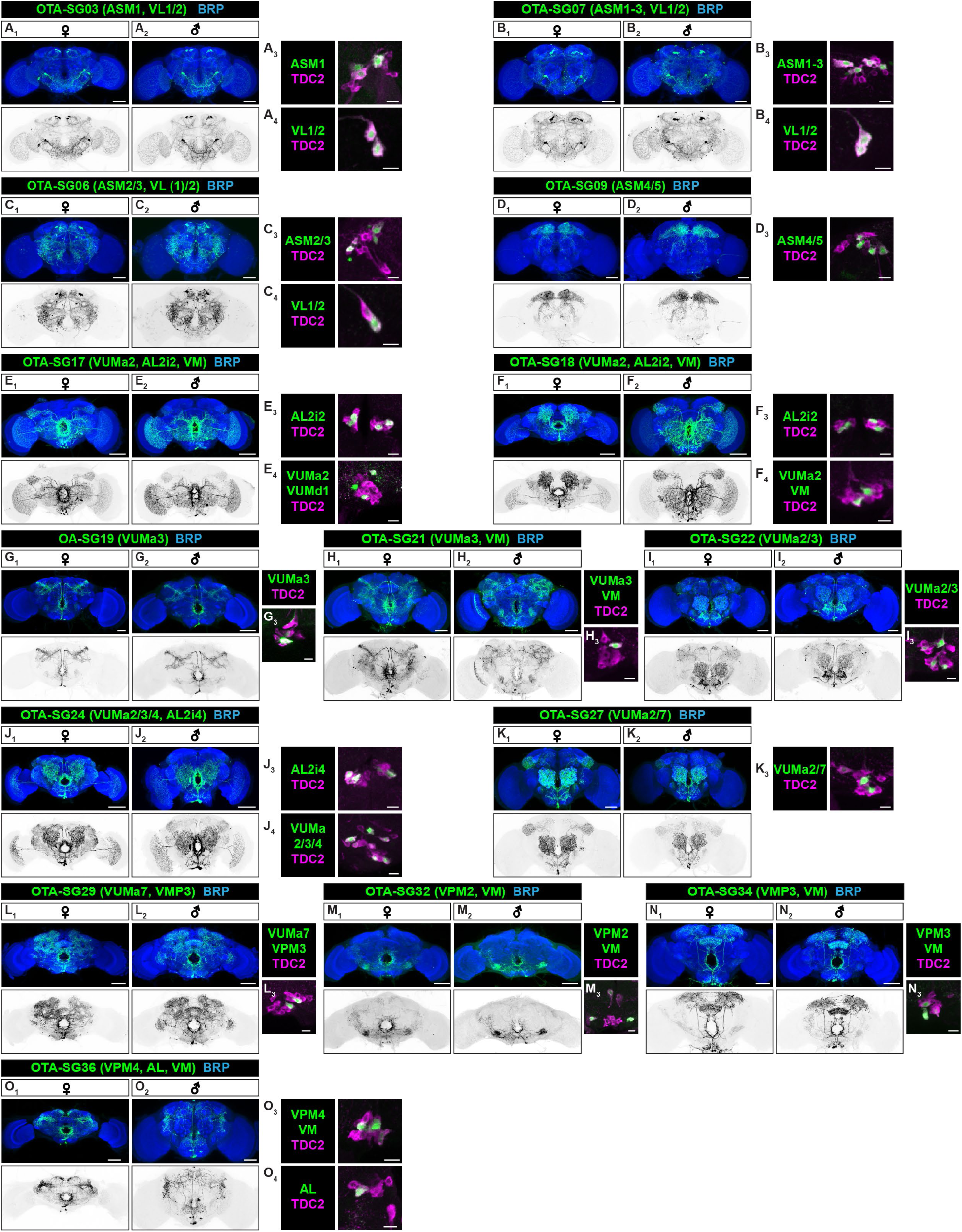
related to Figures 1-4: Additional split-GAL4 lines for OA/TA subtypes. Representative images of female (**A_1_**-**O_1_**) and male (**A_2_**-**O_2_**) brains labeled by split-GAL4 lines (OTA-SG) not described in the main figures. Panels show stacked confocal images, with neuropils (visualized by anti-bruchpilot: top panels, blue) and CsChrimson-tagged tdTomato expressed by the given split-GAL4 line (visualized by anti-dsRed: top panels, green; bottom panels, black). Scale bar = 50 μm. Targeted cell types are indicated along with a split-GAL4 line name. (**A_34,_**, **B_3,4_**, **C_3,4_**, **D_4_**, **E_3,4_**, **F_3,4_, G_3_, H_3_, I_3_, J_3,4_, K_3_, L_3_, M_3_, N_3_, O_3,4_**) Representative zoomed-in images of GFP-expressing OA/TA cell clusters targeted by split-GAL4 line (visualized by cytosolic GFP: green) that are co-labeled by TDC2 immunoreactivity (magenta). Scale bar = 10 μm.

**Supplementary Figure S3.**
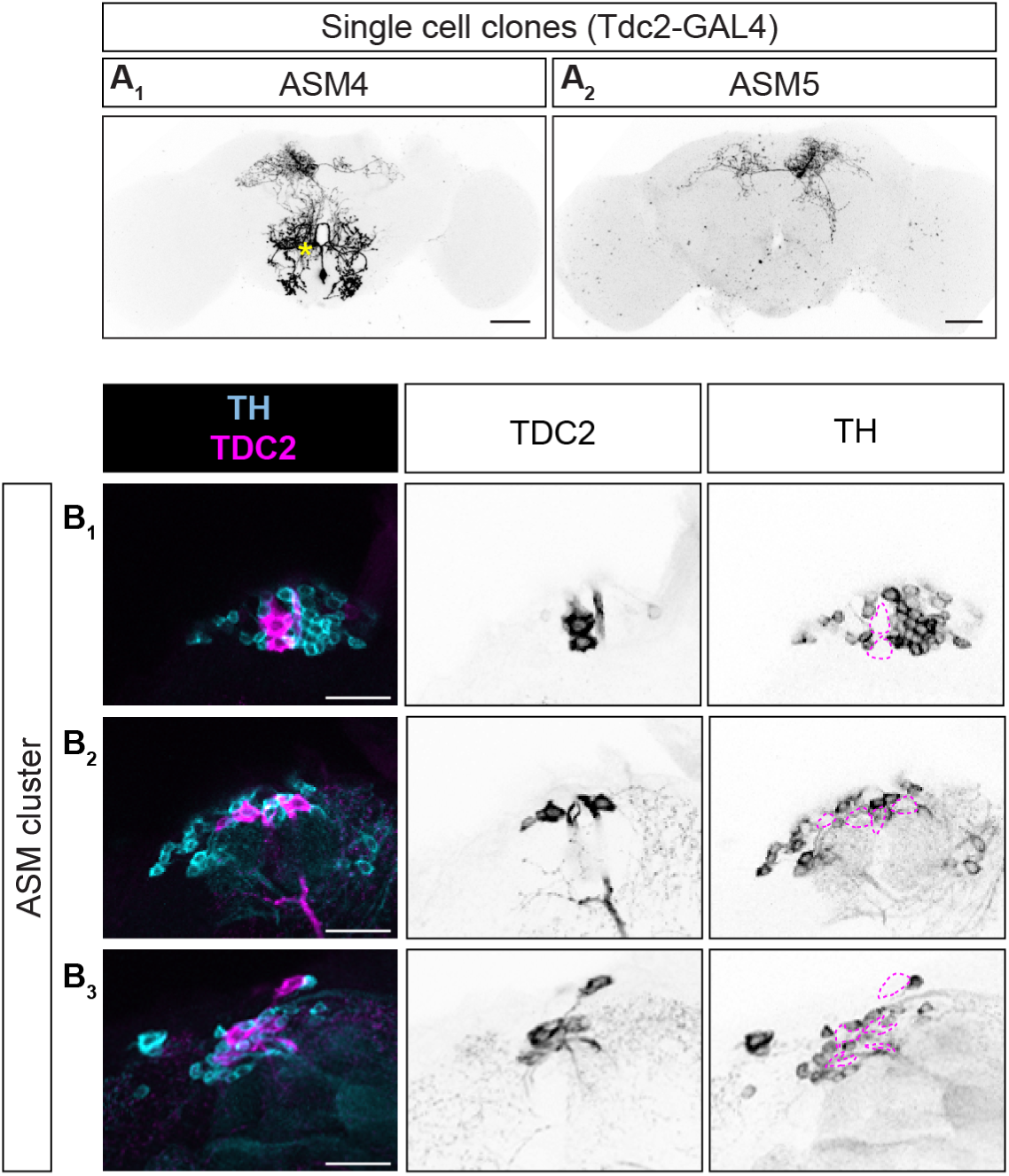
related to Figure 1: Detailed characterization of ASM neurons. (**A**) Single cell clones of ASM4 (**A_1_**) and ASM5 (**A_2_**) from *Tdc2-GAL4* obtained with MCFO technique. A non-ASM4 neuron (VPM1) is indicated in yellow asterisk. Scale bar = 50 μm. (**B**) Three representative confocal image sections from a single sample (**B_1-3_**, from anterior to posterior) of Tdc2-expressing ASM neurons (left panels, magenta; middle panels, black) co-labeled by the antibody against the tyrosine hydroxylase (TH) (left panels, cyan; right panels, black), which is encoded by the *pale* gene in the *Drosophila* genome. No co-localization of TDC2 and TH was observed even though ASM neurons are intermingled with PAM dopaminergic neurons. Scale bar = 20 μm.

**Supplementary Figure S4.**
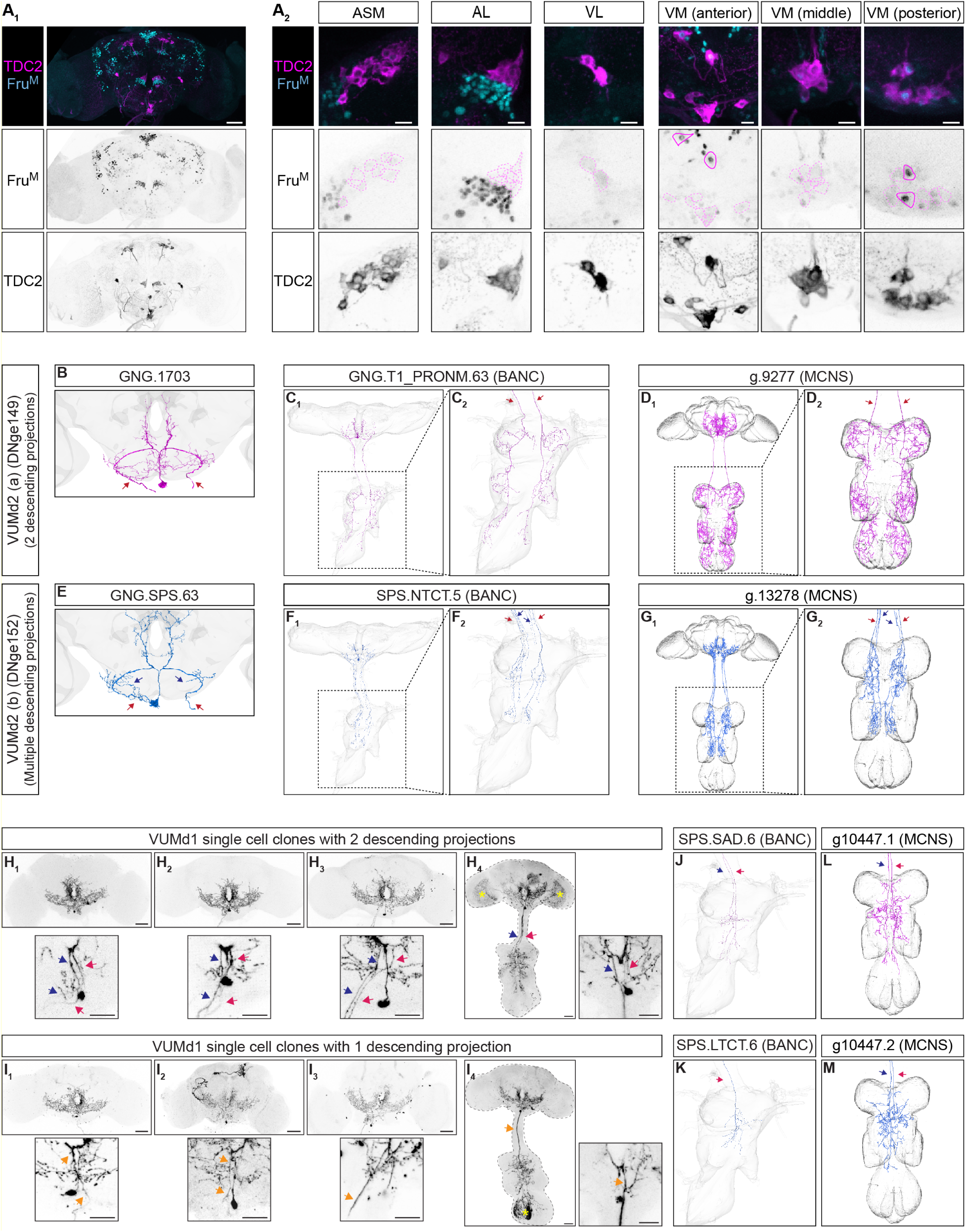
related to Figure 4: Additional characterization of VUMd neurons. (**A**) Representative images of OA/TA neurons in an adult male brain visualized by TDC2 immunoreactivity (top panels, magenta; bottom panels, black) along with immunoreactivity against the male-specific isoform of Fruitless (FruM) (top panels, cyan; middle panels, black), both as a whole-brain z-stack of a confocal volume image (**A_1_**; Scale bar = 50 μm), and as zoomed-in images of TDC2-positive neuronal clusters (**A_2_**; Scale bar = 10 μm) FruM-positive OA/TA neurons are present only in the VM cluster (see VM-anterior and VM-posterior panels). (**B-G**) Neurons annotated as VUMd2 in EM volumes are split into two cell types. DNge149 in FAFB (**B**), BANC (**C**), and MCNS (**D**) have 2 parallel descending projections to VNC that pass the neck connective laterally (**C_2_, D_2_**). DNge152, by contrast, have multiple, fine descending projections (**E-G**). Unlike DNge149 (**C_2_, D_2_**), DNge152 does not innervate to the 3^rd^ prothoracic neuromere (**F_2_, G_2_**). (**H, I)** Single cell clones from *Tdc2-GAL4* (**H_1,2_**, **I_1_**) and *OTA-SG37* (**H_3,4_**, **I_2-4_**) obtained with MCFO showed two types of descending projections. Some have two descending projections (**H**, blue and magenta arrows), while others have just one projection (**I**, orange arrows), that originate from the esophagus foramen. Non-VUMd1 neurons are indicated in yellow asterisks. Scale bar = 50 μm for top panels, 25 μm for bottom panels (zoomed section). (**J-M**) The BANC EM volume has one DNge138 (VUMd1) trace with two descending projection (**J**) and one trace with one descending projection (**K**), while two DNge138 traces in the MCNS EM volume both have two descending projections (**L, M**). Images of H_1_, H_3_, and I_1_ are reproduced from Fig. 4. G_3_, G_1_, and I_2_, respectively.

**Supplementary Figure S5.**
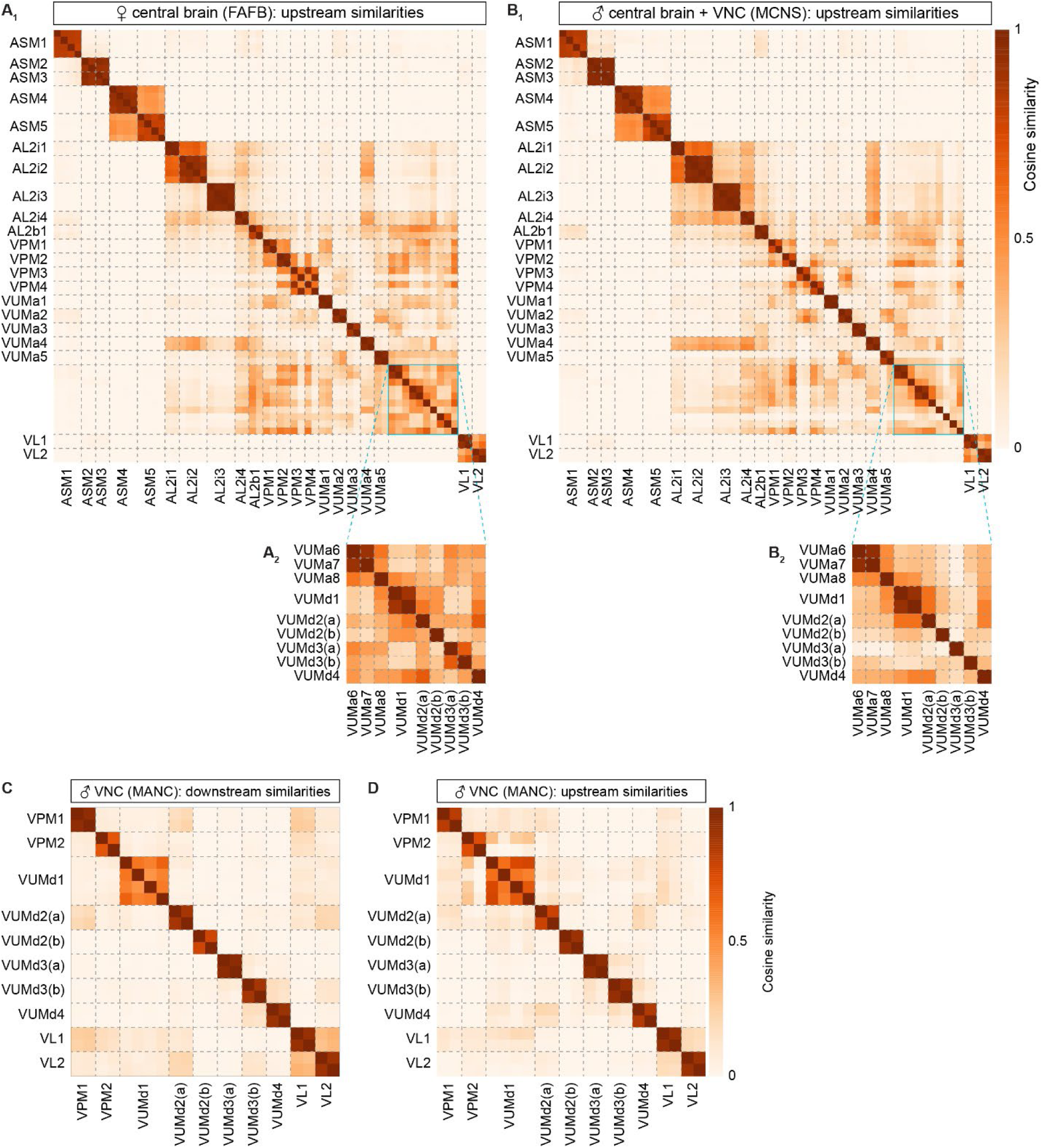
related to Figure 5: Additional connectivity analyses of OA/TA cell types. (**A, B**) Connectivity similarity of OA/TA neurons identified in the EM volumes of the female central brain (FAFB) (**A**) and the male central brain and nerve cord (MCNS) (**B**) based on their upstream neuronal types. Broken lines represent the cell type boundaries. Regions that include a cell type containing only one neuron (cyan squares in **A_1_**, **B_1_**) are enlarged below (**A_2_**, **B_2_**). (**C, D**) Connectivity similarity of OA/TA neurons identified in the EM volumes of the male ventral nerve cord (MANC) based on their downstream (**C**) and upstream (**D**) neuronal types. Broken lines represent the cell type boundaries. VPM1, VPM2, VL1, and VL2 has one corresponding trace per neuron in VNC because these neurons send a unilateral descending projection. VUMd1-5 have two corresponding traces per neuron in VNC because these neurons send largely bilateral descending projections (see also main text).

**Supplementary Figure S6.**
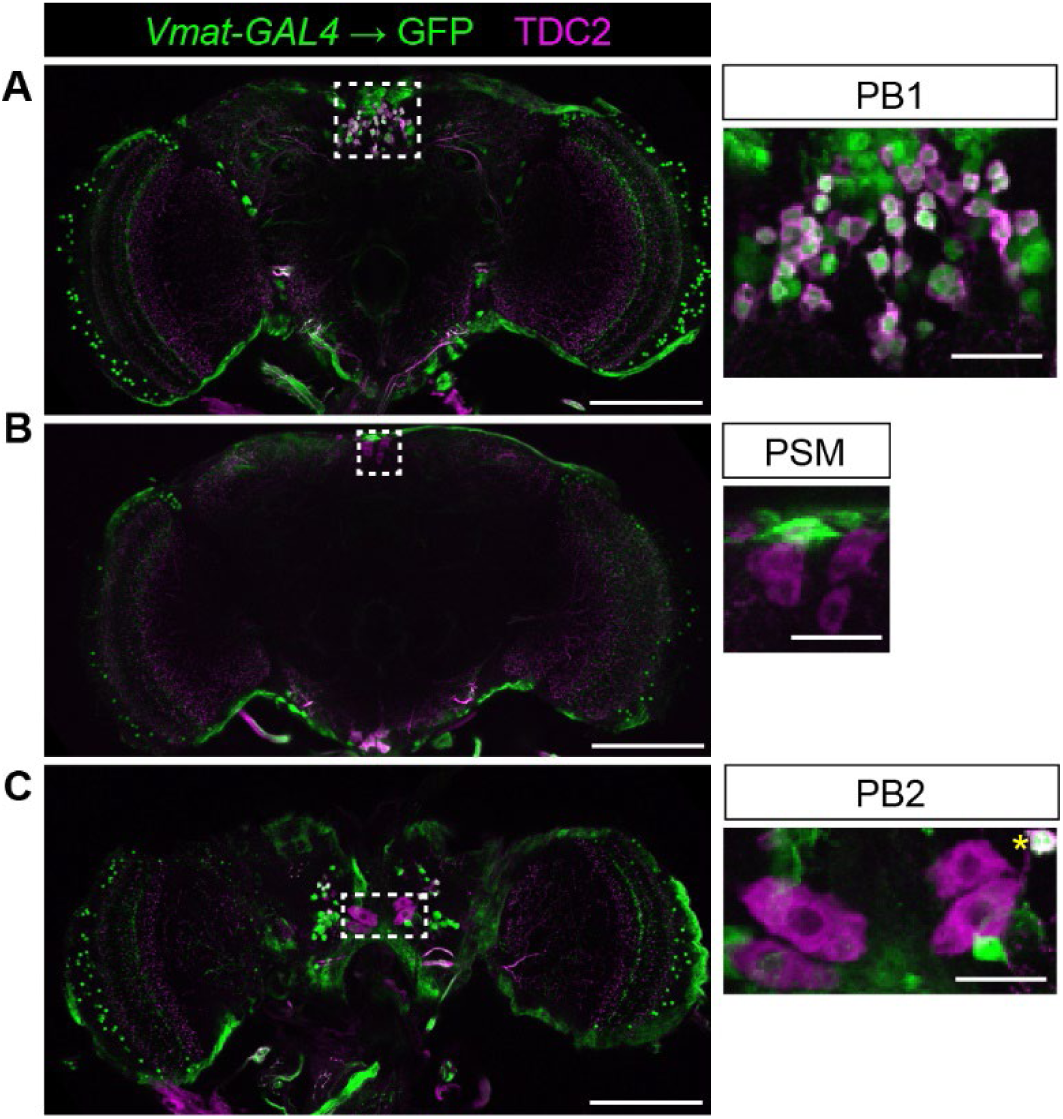
related to Figure 1-4: PB1 cluster expresses *Vmat*. Representative images of male brain with immunohistological visualization of cytosolic GFP (green), expressed under *Vmat-GAL4*, and TDC2 (magenta). (**A**) Tdc2-positive cells in PB1 cluster co-localize with GFP. In contrast, Tdc2-positive cells in PSM (**B**) and (**C**) P2 clusters do not co-localize with GFP. Scale bar = 100 μm for left panels, 20 μm for right panels (zoomed section). An yellow asterisk in the left panel of C indicates a nearby PB1 neuron.

